# Predicting TCR antigen specificity at proteome-scale with synthetic immune cells and machine learning

**DOI:** 10.1101/2025.07.23.666264

**Authors:** Kai-Lin Hong, Beichen Gao, Lucas Stalder, Roy A. Ehling, Marton Horvath, Sarah Wehrle, Juliette L. Forster, Valentin Junet, Jakub Kucharczyk, Sébastien Lalevée, Marie-Charlotte Diringer, Qinmei Yang, Rodrigo Vazquez-Lombardi, Sai T. Reddy

## Abstract

TCR specificity to peptide-HLA antigens is central to immunology, impacting responses in infection, autoimmunity and cancer. Achieving precise recognition while avoiding off-target reactivity is critical for effective immunity and safe therapeutic interventions. Comprehensive, proteome-wide specificity profiling of TCRs is challenging with current methods, which notably lack integrated machine learning for large-scale analysis. Here, we report a synthetic immune cell system coupled with machine learning to enable TCR functional and specificity mapping of peptide-HLA antigens at proteome-scale. Multi-step immunogenomic engineering of synthetic antigen-presenting cells (APCs) was performed to enable stable mono-allelic integration and precise display of peptide antigens and HLA class I from a defined genomic locus, ensuring genomically-encoded antigen presentation. Compatible with a synthetic TCR displaying T cell system, this platform incorporates a fluorescent reporter of cytokine-mediated signaling for real-time activation detection in both synthetic APCs and T cells. We combined this screening with diverse peptide antigen libraries and deep sequencing to train supervised machine learning models. These models were applied to predict TCR specificity to peptide-HLA antigens across the entire human proteome. Experimental validation confirmed novel off-targets for therapeutic TCR candidates, including for a clinically-approved TCR therapeutic. This integrated synthetic immune cell and machine learning approach provides unprecedented proteome-wide peptide-HLA specificity mapping to support the development of safer TCR-based therapies.

**One sentence summary:** We present a synthetic immune cell platform integrated with machine learning that enables prediction of TCR specificity to peptide-HLA antigens at proteome-scale.

## Introduction

T cell receptors (TCRs) are central to adaptive immunity, mediating the recognition of peptide fragments presented by human leukocyte antigen (peptide-HLA, pHLA) on cell surfaces. This specific recognition allows the immune system to distinguish diseased cells, such as those infected by pathogens or transformed by cancer, from healthy tissue. The ability of TCRs to target intracellular antigens, which constitute the vast majority of the proteome, offers a larger target space relative to traditional antibody-based approaches restricted to surface antigens. This expanded target repertoire makes TCR-based therapies a highly promising avenue for treating a wide array of cancers and potentially other diseases driven by intracellular self or foreign antigens (*1*).

Despite this therapeutic promise, a key challenge for TCR-based therapies is the risk of off-target reactivity. Engineered TCRs, often optimized for high-affinity binding to a target pHLA on cancer cells, may recognize structurally similar but distinct pHLA presented on healthy tissues. Such cross-reactivity can lead to severe adverse events, as exemplified in past clinical trials of affinity-enhanced TCR-T cells targeting the cancer-testis antigen MAGE-A3 (*2–5*). These examples highlight a key property of natural TCR selection, which allows for a degree of cross-reactivity for broad pathogen protection (*6*). Enhancing the affinity of TCR drug candidates for their intended targets can exacerbate cross-reactivity potential, thereby increasing the risk of harmful off-target effects. Consequently, ensuring precise on-target recognition while meticulously avoiding off-target interactions is essential for the development of safe and effective TCR-based therapeutics.

Bispecific T cell engagers based on soluble TCR (sTCR) fragments (*7*) and TCR-like antibodies (*8*) that bind pHLA with high affinity represent an emerging therapeutic modality for redirection of polyclonal T cells to tumor targets. The clinical approval of Kimmtrak (Tebentafusp), a sTCR-anti-CD3 bispecific targeting gp100/HLA-A*02:01 for uveal melanoma, has demonstrated the therapeutic potential of sTCR engagers (*9*) and motivated the development of numerous other pHLA-targeting programs. This underscores a growing need for platforms capable of comprehensively assessing the specificity profiles of these drug candidates.

A number of methods exist for profiling TCR specificity. Peptide scanning involves using individual synthetic peptide point mutants of pMHC to profile TCR specificity and predict off-targets, however is limited by only being able to capture a limited part of the combinatorial mutational space. High-throughput methodologies have been developed for profiling TCR specificity encompass binding-based assays, such as yeast surface display (*10*) and pMHC multimer-barcode labeling (*11*). Functional screening platforms using mammalian cells developed in recent years utilize diverse readouts, including trogocytosis (*12*), engineered MHC receptors with synthetic signaling domains (*13*, *14*), detection of T cell-secreted proteases (*15–17*) or cytokines (*18*), and synthetic reporter protein circuits within APCs (*19*, *20*). The use of genome-wide libraries, in particular, has been demonstrated as an effective approach to identify potential off-targets as well as for TCR de-orphaning (*17*). While the functional reporter aspects of these systems are useful to screen specificity of TCRs in the micromolar affinity range (e.g., for TCR-T cell therapy candidates), affinity-enhanced TCRs with nanomolar-to-picomolar affinities are not suitable for this type of screens due to high background activation of T cells. Furthermore, mammalian display systems are limited by the diversity of libraries that can be feasibly screened, with an upper limit of approximately 10^6^ library members. Crucially, existing screening platforms have not yet integrated machine learning (ML) for robust, proteome-scale prediction of TCR specificity, a key step for comprehensive off-target risk evaluation.

Here, we describe an integrated synthetic immune cell system coupled with ML to enable TCR specificity predictions to peptide-HLA antigens at proteome-scale. To achieve this, we first engineered a synthetic antigen presenting cell (APC) through multi-step, targeted immunogenomic engineering to achieve stable, mono-allelic, genomically-encoded display of diverse peptide antigens and HLA class I molecules from a defined genomic locus. In this manner, the system leverages natural peptide antigen presentation to enable accurate presentation of pHLA libraries on the surface of mammalian cells. The system further incorporates a fluorescent reporter of cytokine-mediated signaling, enabling functional validation of predicted off-targets using the same cell line. We used our system to screen two affinity-enhanced sTCR engagers, including the clinically approved drug Kimmtrak, against diverse peptide antigen libraries, followed by deep sequencing to generate datasets used to train supervised ML models. These models are used to predict TCR specificity to pHLA antigens across the entire human proteome. ML predictions and experimental validation confirmed novel off-target peptides, including peptides with a large edit distance to the target epitope, thus demonstrating the platform’s capacity to support the development of safer TCR-based therapies

## RESULTS

### Development of a synthetic antigen-presenting cell platform through multi-step genome engineering

We developed the antigen-presenting cell detecting cytokine (ACDC) platform through stepwise CRISPR-Cas9 genome engineering. ACDC cells utilize a fluorescent cytokine reporter and genomically-encoded antigen presentation to enable high-throughput functional screening of TCR and peptide–HLA-I (pHLA) interactions. The ACDC system was established by engineering a CRISPR-edited monoclonal derivative of HEK293 cells to constitutively express Cas9-GFP from the CCR5 locus and a STAT5-driven mRuby2 reporter of IL-2 signaling (Fig. 1A and sFig. 1). This configuration enables fluorescent detection of antigen-specific TCR activation via paracrine IL-2 signaling from co-cultured T cells. To reduce background antigen presentation, we performed a pan-HLA class I knockout using a conserved exon 4-targeting guide RNA (gRNA), resulting in complete ablation of endogenous HLA-A, -B, and -C surface expression (Fig. 1, B and sFig. 1). To ensure mono-allelic insertion of peptide–HLA constructs, one allele of the CCR5-integrated GFP was first replaced with BFP, creating two phenotypically traceable landing pads (Fig. 1, C and D). With the landing pads in place, we performed mono-allelic insertion of a HLA-A*02:01 transgene into the BFP-containing CCR5 allele. Successful knock-in was confirmed by flow cytometry (Fig. 1, D and E) and genomic PCR (Fig. 1F), resulting in a clonal ACDC-HLA cell line that stably expressed transgenic HLA-I and retained IL-2 responsiveness via STAT5-mRuby2 activation. Using the same genome engineering approach, additional ACDC-HLA cell lines expressing 18 distinct HLA class I alleles were generated, thus confirming a broad applicability of the platform for multiple HLA-I backgrounds (sFig. 2).

**Fig. 1.**
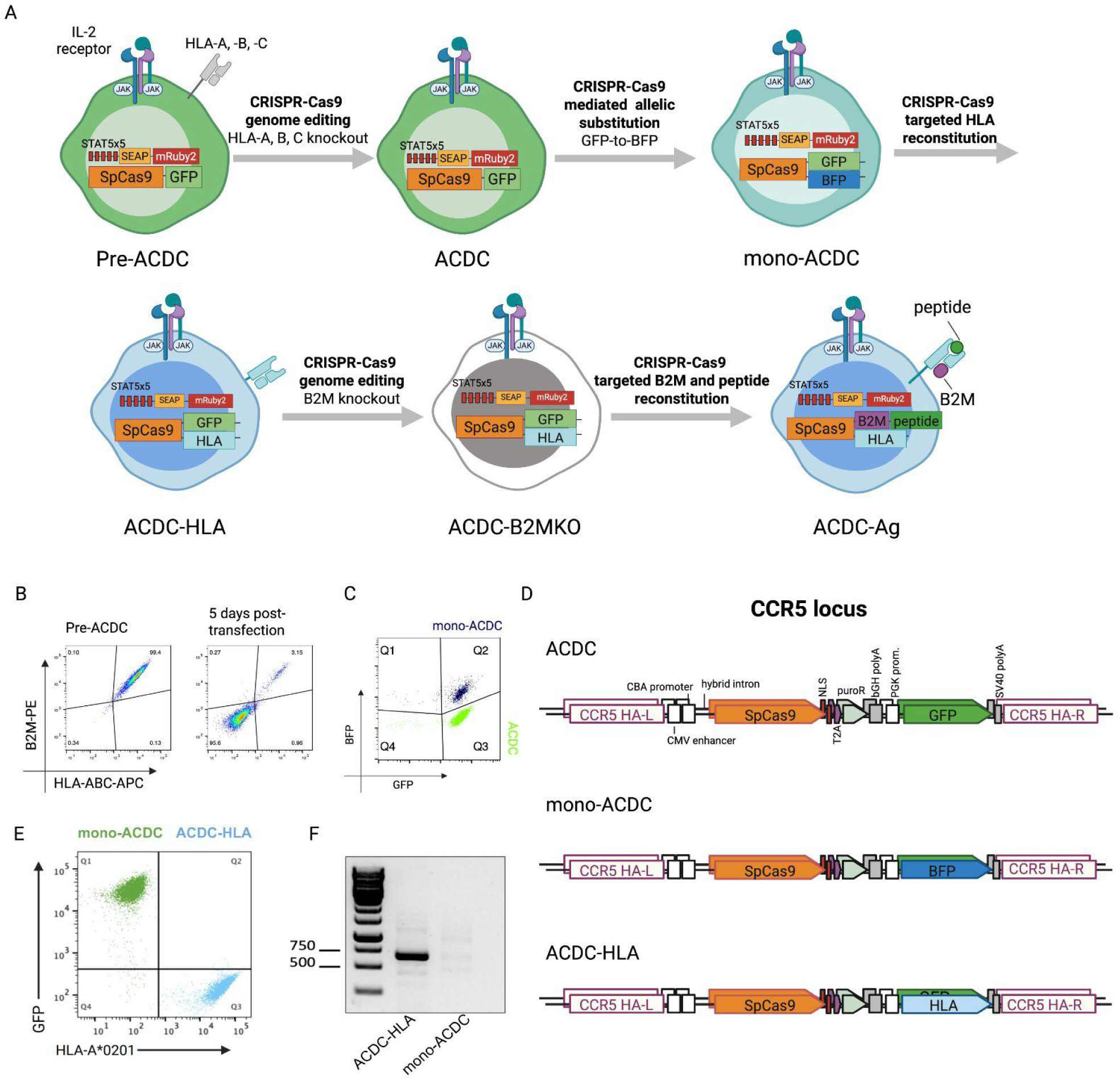
Development of the ACDC synthetic antigen-presenting cell platform through multi-step CRISPR-targeted genome editing. (A) Schematic overview of stepwise CRISPR-Cas9 genome engineering used to generate ACDC and its derivatives. (B) Flow cytometry plots of Pre-ACDC and ACDC cells showing efficient knockout of HLA-A, B, C expression following CRISPR targeting. (C) Representative flow cytometry plot confirming successful mono-allelic replacement of GFP with BFP in mono-ACDC cells. (D) Schematic maps of the engineered CCR5 locus in ACDC, mono-ACDC, and ACDC-HLA cells. (E) Flow cytometry plots validating mono-allelic HLA-A*02:01 knock-in into GFP-positive CCR5 allele. (F) Genomic PCR confirming integration of HLA-A*02:01 into the CCR5 locus.

**Fig. 2.**
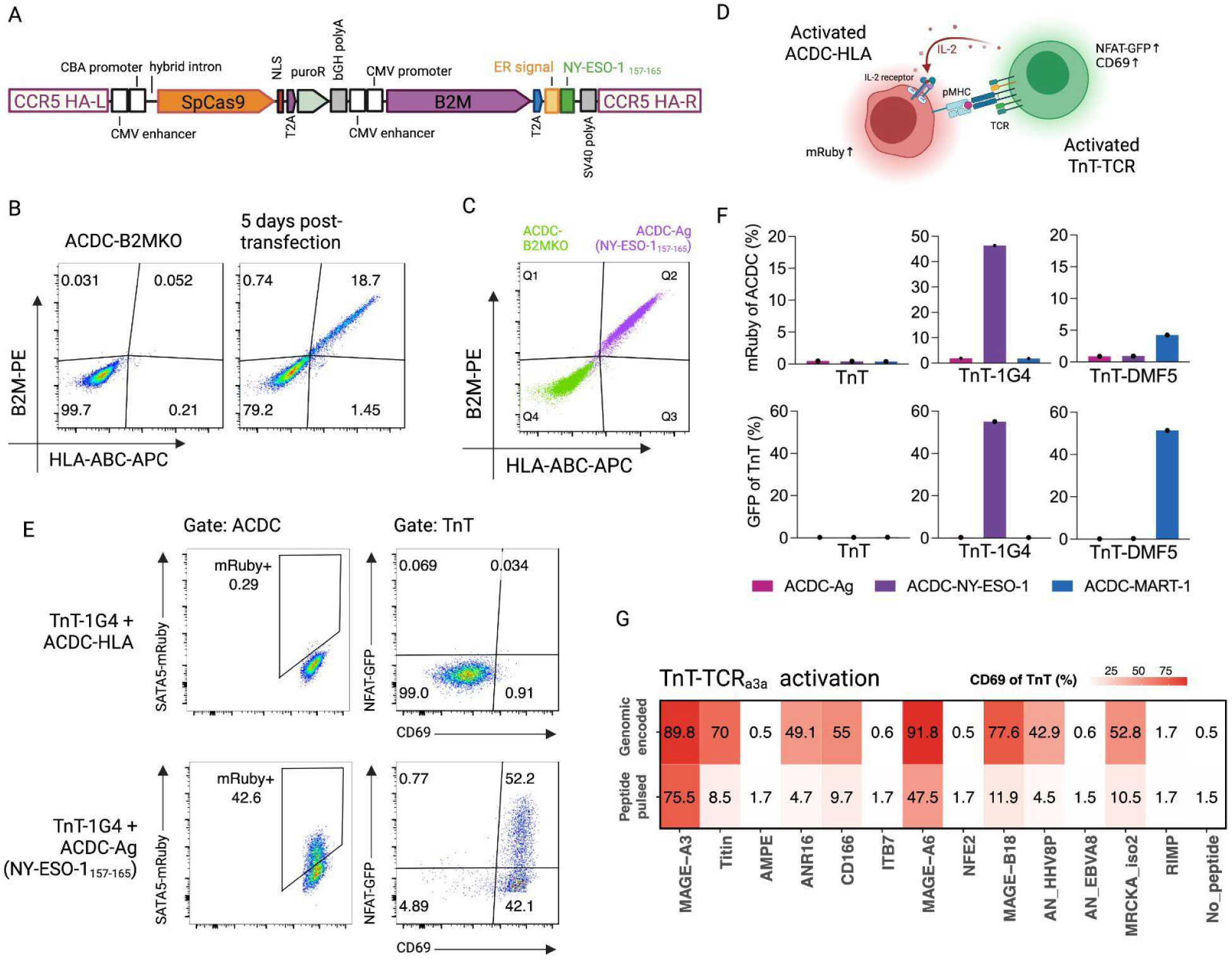
Functional validation of the ACDC-Ag platform for screening TCR specificity and cross-reactivity. (A) Schematic of the HDR template used for antigen reconstitution, encoding a B2M and NY-ESO-1_157–165_ separated by a T2A, preceded by an ER signal sequence to promote proper HLA loading. (B) Flow cytometry analysis of ACDC-B2MKO cells five days post-transfection shows efficient reconstitution of B2M and peptide antigen expression following CRISPR-mediated HDR. (C) Enrichment of B2M–peptide positive cells by FACS, generating clonal ACDC-Ag (NY-ESO-1_157–165_) cells. (D) Schematic of the TnT/ACDC co-culture system. Antigen-specific activation of TnT-TCR cells is measured via NFAT-GFP and CD69, while ACDC cells that express cognate pHLA are detected through STAT5-mRuby2 expression upon IL-2 sensing from activated TnT-TCR cells. (E) Flow cytometry analysis of NFAT-GFP, CD69, and mRuby2 signals following co-culture of TnT-TCR_1G4_ cells with ACDC-HLA or ACDC-Ag cells presenting NY-ESO-1_157–165_. (F) Quantification of IL-2 signaling (mRuby2) in ACDC cells and NFAT-GFP in TnT cells, co-cultured with ACDC-Ag cells expressing NY-ESO-1 or MART-1 peptides, confirming activation is restricted to cognate TCR–pHLA interactions. (G) Heatmap of CD69 expression in TnT-TCR_a3a_ cells co-cultured with ACDC-Ag or peptide-pulsed APCs presenting 13 candidate antigens. Comparable activation confirms the fidelity of genomic antigen presentation by ACDC cells.

To enable genomic integration of constructs encoding peptide antigens and provide a phenotypic marker for successful knock-in events, we next disrupted endogenous β2-microglobulin (B2M), an essential component of HLA-I surface expression (sFigs. 1 and 3A). The resulting ACDC-B2MKO cells exhibited complete loss of B2M and HLA-ABC surface expression, establishing a clean background reconstitution with transgenic pHLA. To reconstitute the ACDC-B2MKO cells with a desired antigen, we designed a homology-directed repair (HDR) template encoding B2M fused to the NY-ESO-1_157-165_ (SLLMWITQC) peptide antigen via a self-processing T2A element. The construct was designed for expression under the control of a CMV promoter and included an N-terminal endoplasmic reticulum (ER) signal sequence upstream of the peptide to ensure its trafficking into the ER for loading onto HLA-I molecules (Fig. 2A). Five days post-transfection, approximately 18% of cells exhibited restored surface expression of HLA-B2M complexes, as assessed by flow cytometry (Fig. 2B). Following FACS enrichment, we established a clonal ACDC-Ag population capable of presenting the genome-encoded peptide antigen (Fig. 2C). Applying the same approach, we generated additional ACDC-Ag cell lines expressing HLA-A*01:01 and a panel of 12 different peptide antigens, achieving HDR efficiencies ranging from 7% to 31% (sFig. 3B).

**Fig. 3.**
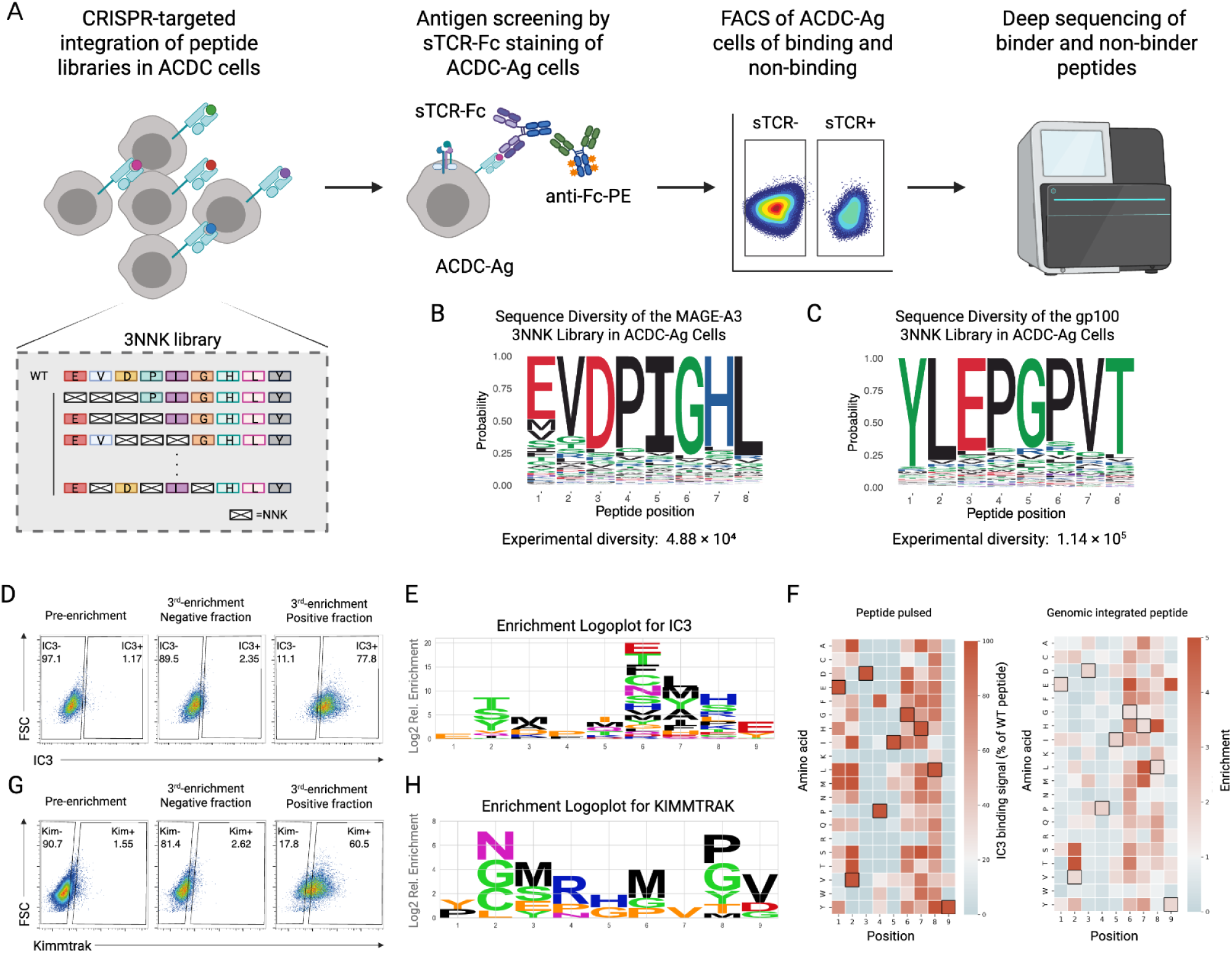
High-throughput screening of peptide–HLA libraries reveals distinct cross-reactivity profiles of soluble TCRs. (A) Schematic overview of the ACDC-based screening pipeline: 3NNK peptide libraries are genomically integrated into ACDC-Ag cells, screened via soluble TCR-Fc (sTCR-Fc) binding, followed by FACS and deep sequencing of binding and non-binding populations. (B) Sequence logo plots of MAGE-A3 3NNK libraries show diverse amino acid coverage across randomized positions. (C) Sequence logo plots of gp100 3NNK libraries show diverse amino acid coverage across randomized positions. (D) Representative FACS plots showing enrichment of TCR_MAG-IC3_-Fc binders after three rounds of selection. (E) Heatmaps comparing cross-reactivity profiles of TCR_MAG-IC3_ from peptide-pulsed (left) and genomically encoded (right) peptide libraries, showing comparable patterns. (F) Enrichment logoplots highlight cross-reactive amino acid profiles of TCR_MAG-IC3_. (G) FACS analysis of Kimmtrak–Fc binding enrichment across three selection rounds. (H) Enrichment logoplot for Kimmtrak showing a more restricted binding motif.

To validate their functionality, we co-cultured ACDC-Ag cells presenting genomically-encoded pHLA targets with an engineered human T cell line previously established by our group (TnT cells) (*21*). TnT cells can be readily reconstituted with individual TCRs or TCR libraries and employed for functional screening through a fluorescent reporter of T cell activation (NFAT-GFP) or CD69 surface expression following co-culture with APCs and cognate pHLA (Fig. 2D). TnT cells were generated expressing TCR_1G4_ (specific for NY-ESO-1_157–165_) and TCR_DMF5_ (specific for MART-1_27–35_), both recognising HLA-A*02:01-restricted peptide antigens. In addition to TnT-TCR cell activation, we assessed STAT5-mRuby2 fluorescence in ACDC-Ag cells as a reporter of IL-2 secretion from activated TnT-TCR cells. Co-culture of TnT-TCR_1G4_ cells with ACDC-Ag cells expressing NY-ESO-1_157–165_ induced robust activation across all markers. In contrast, co-culture with antigen-negative ACDC-HLA cells yielded no detectable NFAT-GFP, CD69, or mRuby2 expression (Fig. 2E). To further confirm activation was specific to TCR-pHLA recognition, we performed co-cultures of TnT, TnT-TCR_1G4_, and TnT-TCR_DMF5_ cells with ACDC-Ag cells presenting NY-ESO-1_157–165_, MART-1_27–35_, or no antigen. NFAT-GFP expression in TnT cells, as well as mRuby2 fluorescence in ACDC-Ag cells, was strictly limited to cognate TCR–pHLA pairings, with no activation observed in any mismatched or antigen-negative controls (Figs. 2E and 2F). Additional validation was conducted using TnT cells expressing the affinity-enhanced TCR_a3a_, specific for MAGE-A3_168-176_ (*3*), co-cultured with ACDC-Ag cells presenting HLA-A*01:01 and 13 genomically-encoded antigens (Fig. 2G and sFig. 3C). The observed activation profiles confirm that genomically-encoded peptide antigens in the ACDC system enables functional presentation and TCR engagement, yielding responses comparable to those induced by conventional peptide-pulsing of APCs (*21*). In addition, successful activation with a full-length MAGE-A3 construct confirmed that ACDC cells can process and present endogenously expressed antigens through the native antigen presentation pathway for recognition by antigen-specific TCRs (sFig. 4).

**Fig. 4.**
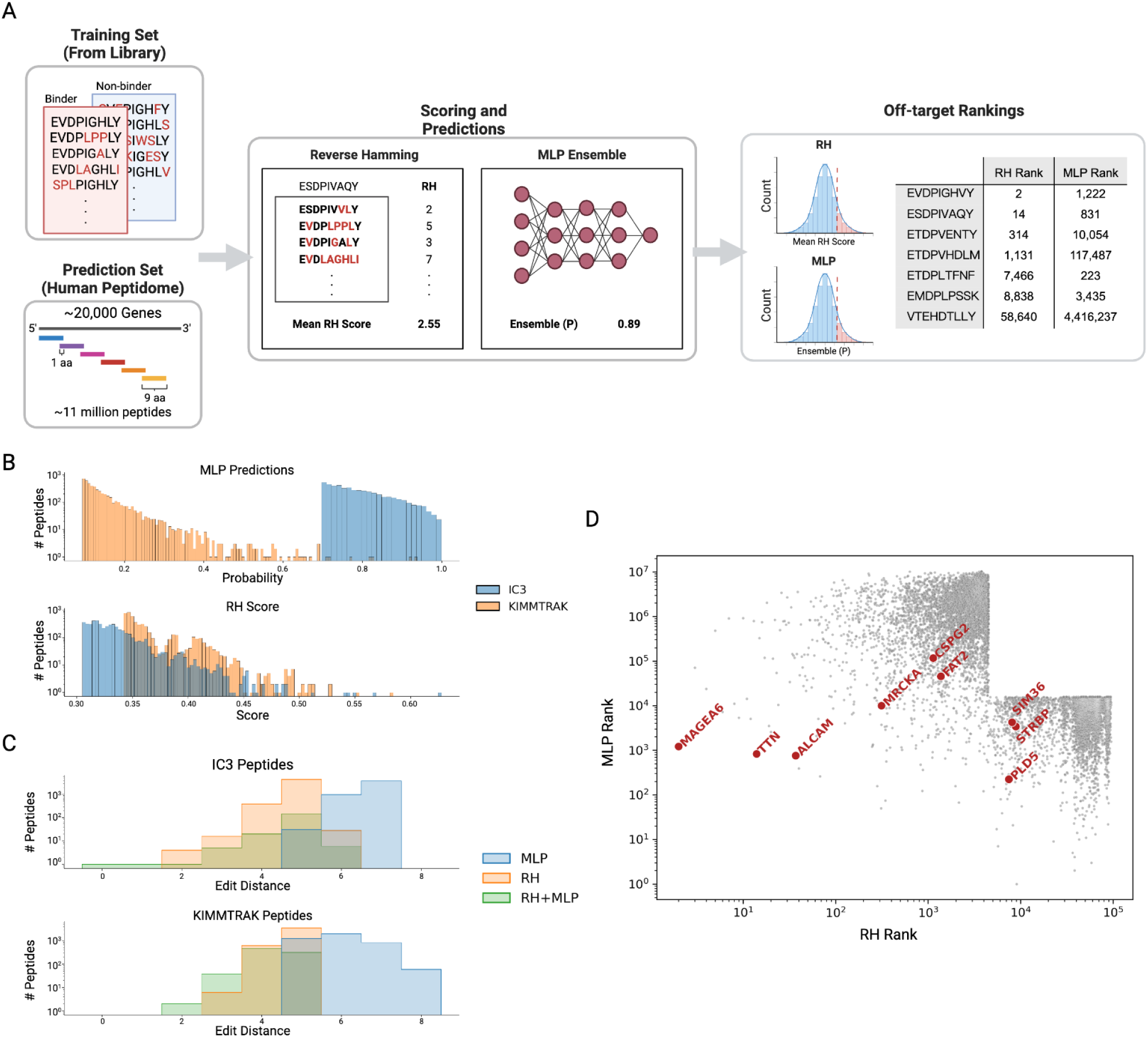
Computational predictions of off-target pHLA antigens across the human proteome. (A) Schematic of the prediction workflow. (B) Distribution of MLP binding probabilities (top) and RH scores (bottom) for the top 5,000 predicted peptides for TCR_MAG-IC3_ (orange) and Kimmtrak (blue). (C) Distribution of edit distances of predicted off-target peptides from the wild-type epitope for IC3 (top) and Kimmtrak (bottom), as predicted by RH (green) and MLP (orange). (D) Scatter plot of RH and MLP rank for the top 5,000 predicted peptides. Known off-targets of TCR_MAG-IC3_ are highlighted in red.

### Screening and sequencing of peptide-HLA libraries against TCR therapeutic candidates

Following engineering and validation of the ACDC platform, we proceeded to screen the cross-reactivity of soluble TCR (sTCR) drug candidates at high-throughput. As a proof-of-concept, we first tested TCR_MAG-IC3_, an affinity-enhanced sTCR targeting MAGE-A3 in the context of HLA-A*01:01, which was originally developed as a drug candidate but presumably discontinued due to off-target reactivity against Titin (*22*), making it an ideal model for validating our platform. To demonstrate broader applicability, we also screened Kimmtrak (tebentafusp), an FDA-approved sTCR targeting gp100 in the context of HLA-A*02:01 (*23*), which has shown clinical efficacy with no reported off-targets to date.

To interrogate the specificity profiles of these sTCRs, we designed peptide mutagenesis libraries that introduced substantial sequence variability while preserving the structural framework of the wild-type (WT) epitope. For TCR_MAG-IC3_ screening, the peptide library was designed to introduce up to three amino acid substitutions by placing degenerate NNK codons (N = A, G, C, or T; K = G or T) at all possible combinations of positions within the WT peptide, resulting in a theoretical diversity of 5.89 × 10⁵ (Fig. 3A). This 3NNK library was generated via Golden Gate cloning and integrated into ACDC-B2MKO (HLA-A*01:01) cells. FACS enrichment for HLA-B2M surface expression followed by deep sequencing confirmed expected amino acid frequencies across randomized positions (Fig. 3B and table S1). ACDC-Ag cells expressing the MAGE-A3 library were subjected to pooled functional screening using soluble TCR_MAG-IC3_-Fc binding assays, followed by three rounds of both positive and negative selection (Fig. 3D). Deep sequencing identified 7,173 unique binders and 17,020 non-binders, providing a high-resolution dataset for modeling TCR cross-reactivity. Sequencing read enrichment analysis (Fig. 3E) revealed broad amino acid tolerance across multiple antigen positions, suggesting a permissive binding profile and high cross-reactivity potential of TCR_MAG-IC3_. As an internal control, we benchmarked read frequency enrichment data of single mutants (i.e., 1NNK) against peptide pulsing co-culture results, which confirmed that genome-encoded peptide variants closely recapitulate peptide pulsed variants for sTCR binding (Fig. 3F).

For Kimmtrak screening, we applied the 3NNK design strategy to the gp100 peptide, integrating the resulting library into ACDC-B2MKO (HLA-A*02:01) cells. As before, the library was validated through FACS enrichment of HLA-B2M surface expression and deep sequencing to confirm expected amino acid frequencies (Fig. 3C and table S2). ACDC-Ag cells expressing 3NNK gp100 library variants were screened with soluble Kimmtrak-Fc through three rounds of positive and negative enrichment (Fig. 3G). The resulting enrichment logo plots from the 1NNK subset (Fig. 3H) revealed a narrow profile of amino acid preferences across key positions, indicating a higher degree of target specificity compared to TCR_MAG-IC3_ .

### Proteome-scale screening for off-target peptide antigens by similarity-based scoring and machine learning

To gain deeper insights into TCR specificity profiles and predict their potential off-targets, we analysed the deep sequencing enrichment data using two complementary computational approaches. The first approach is a similarity-based scoring method that involves calculating the reverse hamming (RH) distances of 11 million human 9-mer peptides contained in the UniProt Human Proteome database to the enriched binder set of synthetic peptide antigens identified from our datasets. This method has been used previously to identify target peptides of orphan TCRs (*24*). Notably, however, RH distances inherently bias prediction scores towards peptides with lower mutational distances from the parental antigen.

As our library is limited to peptides with up to three amino acid substitutions from the parental peptide, we additionally employed ML models, which have the potential to generalise to the mutational sequence space beyond our library. We tested a baseline suite of ML classifier models, including logistic regression, XGBoost, and multilayer perceptrons (MLPs), which predict the binding probability of a peptide antigen to a soluble TCR based on its amino acid sequence. Among these, XGBoost achieved the highest Matthews correlation coefficient (MCC), indicating strong overall performance. However, the baseline MLPs demonstrated superior recall, an essential feature for minimizing false negatives in off-target risk assessment (Fig. 4A and table S3). The high performance of XGBoost models suggested ensemble modelling may improve overall performance, thus, we trained an ensemble of 200 MLP models, which showed the best performance on the held out test set, exceeding the performance of XGBoost (table S3). We used this as our final architecture, and trained a final ensemble with 200 MLP models based on full read frequency enrichment data, and used them to predict TCR_MAG-IC3_ binding across the 11 million human 9-mer peptides (table S4). Next, we examined the predicted binding probabilities of the top-ranked 5,000 peptide antigens as a subset as potential off-targets of TCR_MAG-IC3_ (P > 0.5) (Fig. 4B). These predictions were compared against the top ranked 5,000 peptide antigens by RH scores. Despite limited overlap among the top 5,000 candidates from each method (sFig. 5A), both successfully recovered known off-targets of TCR_MAG-IC3_ such as Titin and MAGE-A6 (Fig. 4D and table S5). Notably, the MLP model ensemble uniquely identified off-target peptides from PLD5, STRBP and STIM36 off-targets with an edit distance of six from the wild-type MAGE-A3 peptide, highlighting an enhanced sensitivity of ML for detecting cryptic cross-reactivity (Fig. 4D).

**Fig. 5.**
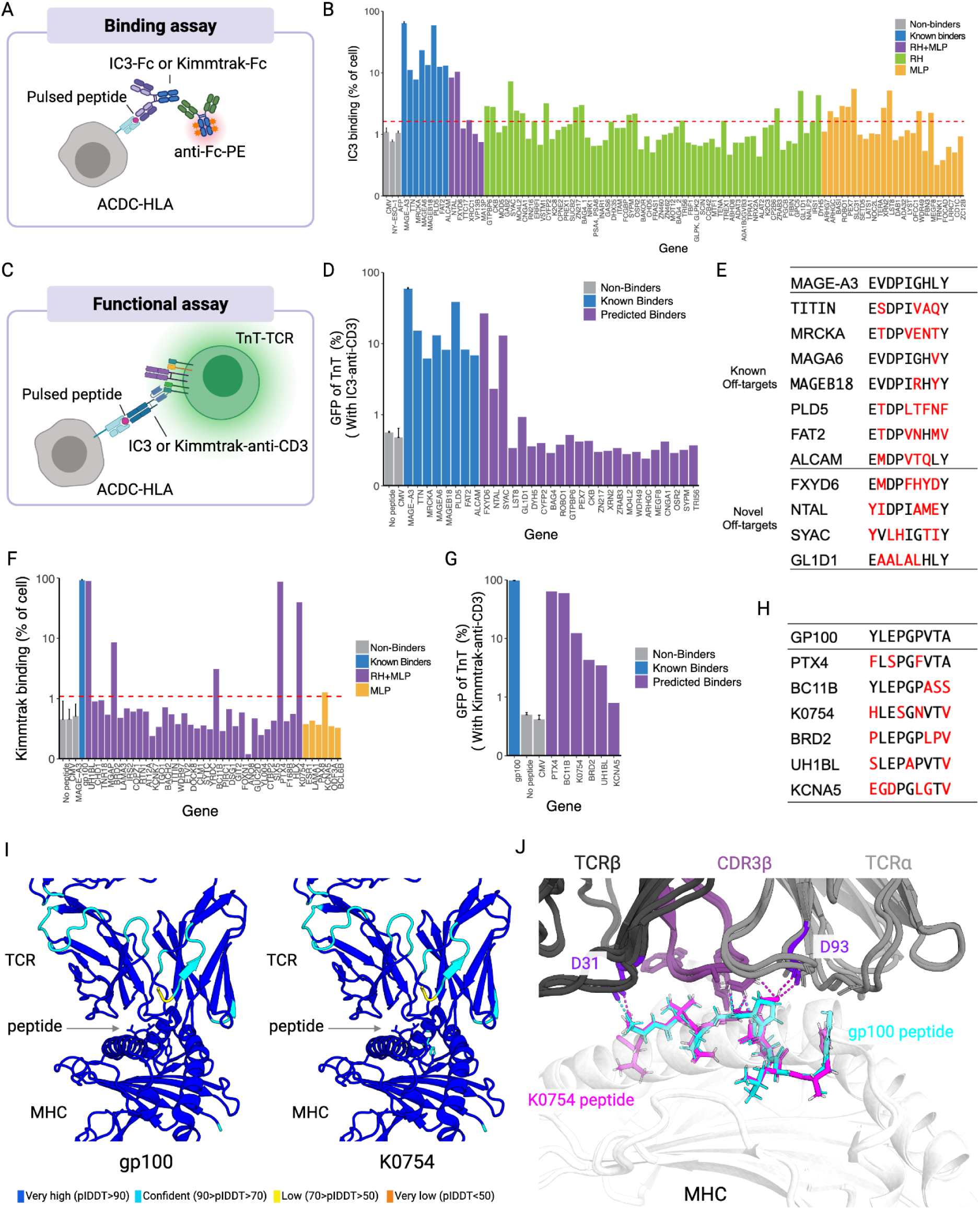
Experimental validation and structural analysis of off-target peptide antigens for TCR_MAG-IC3_ and Kimmtrak. (A) Schematic of the sTCR-Fc binding assay in ACDC-HLA cells pulsed with individual peptides. (B) Quantification of TCR_MAG-IC3_-Fc binding to 30 candidate peptides presented on ACDC-HLA cells. The red dashed line represents a threshold set at three standard deviations above the mean binding of the negative control peptide (CMVpp50). (C) Schematic of the T cell functional activation assay via sTCR-anti-CD3 bispecifics and ACDC-HLA cells pulsed with individual peptides. (D) NFAT-GFP expression in TnT-TCR cells co-cultured with peptide-pulsed ACDC-HLA cells in the presence of TCR_MAG-IC3_-anti-CD3, confirming functional off-target activation. (E) Sequence of known and novel TCR_MAG-IC3_ off-targets, with mutated residues shown in red. (F) Quantification of Kimmtrak-Fc binding to 40 candidate peptides presented on ACDC-HLA cells. The red dashed line represents a threshold set at three standard *deviations above the mean binding of the negative control peptide (CMVpp50*). (G) NFAT-GFP expression in TnT-TCR cells co-cultured with peptide-pulsed ACDC-HLA cells in the presence of Kimmtrak-anti-CD3, confirming functional off-target activation. (I) Structural models of Kimmtrak bound to gp100 and K0754 peptide–HLA-A*02:01 complexes, predicted by AlphaFold3. Color scale indicates pIDDT scores per residue. (J) Overlay of the Kimmtrak–gp100–HLA-A*02:01 crystal structure (PDB: 7PHR) and the AlphaFold3-predicted Kimmtrak–K0754–HLA-A*02:01 complex. TCRβ and TCRα chains are shown in dark and light grey, respectively; the CDR3β loop is highlighted in purple. Peptides are shown as sticks, with gp100 in cyan and K0754 in magenta. HLA-A*02:01 is in white.

We applied the same RH and ML framework to the Kimmtrak enriched-peptide dataset to enable prospective cross-reactivity profiling of an FDA-approved sTCR therapeutic (table S4). In contrast to TCR_MAG-IC3_, MLP probabilities of the top 5,000 peptides had very few instances exceeding a probability of binding above 0.5 (Fig. 4B), suggesting a limited number of potential off-targets. Comparison of RH and MLP scoring again revealed limited overlap in the top 5,000 predicted off-target peptides between the two methods (sFig. 5C), with MLP models tending to predict peptides that differ more substantially from the wild-type gp100 sequence (Fig. 4C, bottom panel). This again highlights a higher sensitivity provided by ML to identify highly distant off-target peptides not captured by similarity-based methods. Interestingly, MLP-predicted off-targets for Kimmtrak exhibited high predicted presentation on HLA-A*02:01, as assessed by NetMHCpan percentile ranks (sFig. 5D), a feature not observed in the TCR_MAG-IC3_ predictions towards HLA-A*01:01 (sFig. 5B).

### Experimental validation of computationally predicted off-target peptide antigens

To experimentally validate predicted TCR_MAG-IC3_ off-targets, we synthesized oligonucleotides encoding the top 5,000 peptides predicted from both RH and MLP models and integrated them into ACDC-Ag cells for pooled functional screening (table S8). After four rounds of TCR_MAG-IC3_-Fc binding enrichment, both libraries reached >50% TCR_MAG-IC3_-positive populations (sFig 6). Deep sequencing revealed a pool of potential binders, from which the 95 highest-ranking candidates (table S9) were selected for individual validation via peptide pulsing (Fig. 5A and B). Of these, 30 antigens demonstrated detectable binding to TCR_MAG-IC3_-Fc. We next screened these 30 antigens for their ability to redirect T cell activation by a bispecific TCR engager molecule. To this end, we produced TCR_MAG-IC3_ as a fusion protein consisting of a TCR_MAG-IC3_ Fab fragment linked to a CD3 agonistic antibody (clone: UCHT1 as scFv antibody fragment) (Fig. 5C and D). ACDC-HLA (A*01:01) cells were pulsed with candidate off-target antigens individually and co-cultured with a TnT-TCR clone expressing endogenous Jurkat TCR, which is not specific to the pulsed antigens, in the presence of soluble TCR_MAG-IC3_-anti-CD3. T cell redirection assays showed that 11 of the 30 peptides triggered activation of TnT-TCR cells (measured by NFAT-GFP expression) via TCR_MAG-IC3_-anti-CD3-mediated engagement (Fig. 5D). This included the known off-target Titin peptide, as well as four newly discovered off-target peptide antigens, FXYD6 (EMDPFHYDY), NTAL (YIDPIAMEY), SYAC (YVLHIGTIY) and GL1D1 (EAALALHLY) (Fig. 5E). Notably, our platform identified off-targets with edit distances of up to six from the MAGE-A3 wild-type epitope. While previously reported off-targets consistently shared identical residues at positions 1, 3, and 4 with the wild-type MAGE-A3 sequence, our newly identified off-targets display a tyrosine (Y) substitution at position 1 (instead of glutamic acid, E), along with diverse mutations at positions 3 and 4.

In contrast to TCR_MAG-IC3_, only 40 peptides were predicted to bind Kimmtrak by the MLP ensembles (P>0.5) (Fig. 4B), 35 of them also among the top 5,000 peptides ranked by RH Score (table S10). Thus, we selected these 40 predicted binders for experimental validation. (table S10). Peptides were synthesized and individually pulsed onto ACDC-HLA (A*02:01) cells, binding by Kimmtrak-Fc revealed six off-target antigens (Fig. 5F). To assess functional relevance, we again performed T cell redirection assays with peptide-pulsed ACDC-HLA (A*02:01) cells and TnT-TCR cells in the presence of a soluble Kimmtrak-anti-CD3 bispecific engager. This revealed that two peptides (BC11B and PTX4) elicited strong T cell activation comparable to gp100, while four additional peptides (K0754, UH1BL, BRD2 and KCNA5) induced lower but detectable activation signals (Fig. 5G). Importantly, these off-targets differed from gp100 by more than three amino acid substitutions, with KCNA5 displaying six mutations, highlighting the platform’s ability to uncover highly divergent cross-reactive epitopes (Fig. 5H).

Kimmtrak has a published crystal structure bound to its cognate antigen, gp100 presented by HLA-A*02:01 (PDB: 7PHR) (*25*). To further examine the structural features of the newly discovered off-targets to Kimmtrak, we performed structural modeling with AlphaFold3 (*26*). For each of the six functionally confirmed off-target peptides (UH1BL, PTX4, K0754, BRD2, BC11B, KCNA5), and the negative control CMVpp50, we generated predictions using 10 independent seeds. For each seed, the top five diffusion-ranked predictions were selected, resulting in 50 structures per complex. Binding to the Kimmtrak TCR was assessed by calculating the average per-atom pLDDT of TCR residues within 5 Å of any peptide residue. All validated peptides and gp100 yielded high-confidence structural models, with per-residue pLDDT scores exceeding 90 across the majority of the complex, particularly at the TCR interface (Fig. 5I and sFig. 7). In contrast, the negative control CMVpp50 peptide resulted in a low-confidence prediction with reduced pLDDT values throughout the peptide-binding region. To investigate the structural basis of cross-reactivity, we overlaid the AlphaFold3-predicted model of the Kimmtrak–K0754 pHLA-A*02:01 complex with the crystal structure of Kimmtrak bound to gp100 pHLA (PDB: 7PHR) (Fig. 5J). Despite the sequence divergence, both peptides exhibited highly similar binding geometries and preserved key interface contacts with the TCR. Specifically, the glutamate at position 3 (E3) and glycine at position 5 (G5) of both gp100 and K0754 engaged the CDR3β loop. Additionally, the conserved threonine at position 7 (T7) in both peptides contacted D31 of the CDR1β loop. Uniquely, K0754 harbored a serine at position 4 (S4), which formed an additional polar contact with both CDR3β and D93 of the CDR3α loop, an interaction not observed in the gp100 complex. These results suggest that conserved and compensatory interactions across both TCR chains underlie the structural mimicry enabling off-target recognition.

## Discussion

Our study establishes a unified experimental-computational workflow that enables proteome-scale interrogation of TCR specificity and cross-reactivity to pHLA antigens. A core innovation of our platform is the use of genome-encoded antigen presentation within the engineered ACDC cell line. Unlike traditional methods, such as yeast display systems that often rely on covalent linking of peptide to HLA molecules (*10*), or mammalian display systems that typically employ transient transfection or viral transduction (*12–20*), our system ensures stable, mono-allelic, genomically-integrated display of peptide antigen libraries. This targeted genomic integration, combined with the natural processing and presentation of pHLA complexes, offers a physiological context for TCR engagement, reducing potential artifacts associated with artificial constructs. This approach uniquely facilitates pooled, high-throughput screening of diverse peptide antigen libraries.

Our workflow couples high-throughput pHLA binding screens with deep sequencing and, critically, ML models to predict off-target specificities across the human proteome. While existing functional mammalian display systems (*17*, *19*) have utilized genome-wide libraries for screening, they have not integrated ML for comprehensive proteome-scale prediction. This integration is a key distinguishing feature of our approach compared to earlier studies. We demonstrate that ML, particularly our ensemble of MLP models, substantially enhances the ability to identify highly distant off-target peptides compared to traditional similarity-based methods such as RH scoring (*27*). For instance, while both RH and MLP models successfully identified known off-targets of TCR_MAG-IC3_, like Titin (four mismatches from the MAGE-A3 epitope), MLP models uniquely identified novel binders, such as PLD5, STRBP, and STIM36, with up to six mismatches. This highlights the capacity of ML to generalize beyond the immediate mutational space of the screened library, uncovering cryptic cross-reactivity patterns that are inaccessible through sequence similarity alone. The experimental validation of these distantly related off-target peptides demonstrates the power of this integrated approach.

The translational relevance of identifying off-targets is paramount for the development of safe and effective TCR-based therapies. As exemplified by the severe adverse events in early clinical trials of affinity-enhanced TCR-T cells targeting MAGE-A3 due to off-target recognition of Titin (*2*, *3*). Our platform identified potential off-targets for the clinically approved bispecific sTCR Kimmtrak (tebentafusp), which highlights its unique sensitivity and capacity to provide a more complete understanding of the specificity profiles of therapeutic TCR drugs in clinical use.

While ML models demonstrated superior recall, essential for minimizing false negatives in off-target risk assessment, we observed that a significant number of predicted off-targets were not experimentally validated, resulting in false positives. This is an inherent characteristic of predictive models, where precision can be traded for sensitivity. However, for drug off-target screening, this conservative approach is often acceptable and even desirable. Prioritizing sensitivity, even at the cost of higher false positives, ensures that a wider range of potential cross-reactive peptides are identified and subsequently experimentally investigated, thereby enhancing TCR drug safety. Further refinements in model architecture, training data diversity, and integration of structural modeling (*26*) may reduce false positives in future iterations.

In summary, this work establishes a scalable and generalizable framework for comprehensive mapping of TCR specificity and cross-reactivity. By uniquely combining genome-engineered immune cells for physiological antigen presentation, high-throughput peptide library screening, and the robust application of ML for proteome-wide specificity assessment, our system provides an unparalleled strategy to increase the throughput, accuracy, and depth of safety screening for soluble TCR engagers and related therapeutic modalities targeting pHLA antigens.

## ACKNOWLEDGMENTS

We thank the ETH Zurich D-BSSE Single Cell facility and the ETH Zurich D-BSSE Genomics facility for their excellent support and assistance throughout this study. This work was supported by ETH Zurich Research Grants.

## AUTHOR CONTRIBUTIONS

K-L.H, B.G., R.V-L. and S.T.R. designed the study; K-L.H, R.A.E, M.H., S.W., J.F., J.K., S.L., M-C.D. and Q.Y. performed experiments; K-L.H, B.G., L.S. and V.J. performed computational analyses; K-L.H, B.G., R.V-L. and S.T.R. wrote the manuscript with input from all authors.

## COMPETING INTERESTS

ETH Zurich has filed for patent protection on the technology described herein, and K-L.H, J.K., R.V.-L., and S.T.R. are named as co-inventors on this patent (WO2024003416A1). S.T.R. holds shares of Alloy Therapeutics, Engimmune Therapeutics, Encelta and Fy Cappa Biologics. S.T.R. is on the scientific advisory board of Alloy Therapeutics, Encelta, Engimmune Therapeutics and Fy Cappa Biologics. RVL holds shares of Engimmune Therapeutics. B.G., R.A.E., M.H., J.F., S.W., V.J., J.K., S.L., M-C.D., Q.Y., R.V-L are employees of Engimmune Therapeutics.

## DATA AND MATERIALS AVAILABILITY

All data are available in the main text or the supplementary materials. Additional data files and code that supports the findings of this study is available from the corresponding authors upon reasonable request. Raw FASTQ files from deep sequencing that support the findings of this study will be uploaded following publication. Original code for the data analysis has been deposited on Github.

## MATERIALS AND METHODS

### Cell lines and cell culture

The HEK-Blue IL-2 cell line was obtained from Invivogen (#hkb-il2-2). The Jurkat leukemia E6-1 T cell line was obtained from the American Type Culture Collection (ATCC) (#TIB152). Engineered TnT cells were cultured in ATCC-modified RPMI-1640 (Thermo Fisher, #A1049101). Engineered ACDC cells and their derivatives were cultured in DMEM medium (ATCC, #30-2002). All the media were supplemented with 10 % FBS (gibco, #16000-044) and 1 % Penicillin-Streptomycin (Gibco, #15140-122). All cell lines were cultured at 37 °C, 5% CO2 in a humidified atmosphere. For prolonged storage, cells were frozen in Bambanker freezing medium (GCLTEC, #BBD01) and stored in liquid nitrogen.

### Polymerase chain reaction (PCR)

PCR for cloning, HDR template generation, genotyping, peptide library construction, and preparation of amplicons for deep sequencing were performed using Q5® Hot Start High-Fidelity 2X Master Mix (NEB, #M0494S) with custom-designed primers (see Supplementary Table 6). Annealing temperatures (x °C) were optimized for each reaction using gradient PCR. The thermal cycling conditions were: initial denaturation at 98 °C for 30 s; 30 cycles of 98 °C for 15 s, x °C for 20 s, and 72 °C for 30 s per kb; followed by a final extension at 72 °C for 2 min.

### Cloning and generation of HDR templates

DNA for gene-encoding regions and homology regions were generated by gene synthesis (Twist Bioscience) or PCR and introduced into desired plasmid backbones via restriction cloning (Supplementary File 1). Targeted knock-in of Cas9/GFP into the CCR5 locus was performed utilizing circular plasmid DNA as the HDR template. HDR templates for all other targeted knock-in experiments were provided as linear double-stranded DNA (dsDNA) generated by PCR. Prior to transfection, PCR products were column-purified using the DNA Clean & Concentrator kit (Zymo, #D4004). For targeted HLA reconstitution of ACDC cells, a double-stranded DNA HDR template was used containing left and right homology arms of 796 and 935 bp, the HLA open reading frame, and an SV40 terminator. HLA cassettes encoding transgenic HLAs were generated by gene synthesis (Twist Bioscience) and cloned into backbone vectors using BamHI and AvrII restriction sites present within the homology arms. Next, HDR templates were generated by PCR using the primer pair RVL-29/RVL-30 and PCR products purified prior to transfection.

### CRISPR-Cas9 genome editing of ACDC cells

Genome editing of ACDC-HLA cells was performed via electroporation using the 4D-Nucleofector device (Lonza) with the SF Cell Line Kit (Lonza, #V4XC-2024). Cells were seeded at a density of 5 × 10⁵ cells/mL one day prior to transfection and cultured for 24 h. Guide RNAs (gRNAs) were assembled by mixing 4 μL of custom Alt-R crRNA (200 μM, IDT) with 4 μL of Alt-R tracrRNA (200 μM, IDT, #1072534), followed by incubation at 95 °C for 5 min and cooling to room temperature. For Cas9-negative HEK-blue cells, 2 μL of assembled gRNA was complexed with 2 μL of recombinant SpCas9 protein (61 μM, IDT, #1081059) at room temperature for at least 10 min to form ribonucleoprotein (RNP) complexes.

Immediately prior to electroporation, cells were washed twice with PBS, and 1 × 10⁶ cells were resuspended in 100 μL of SF buffer. Each reaction included 1.5 μg of HDR template DNA and either 7 μL of gRNA (for Cas9-positive cells) or 4 μL of Cas9 RNP complexes (for Cas9-negative cells). The mixture was transferred into a 1 mL electroporation cuvette and electroporated using program CM130. Post-electroporation, cells were immediately topped up with 1 mL of pre-warmed complete medium and allowed to recover for 10 min before transfer to a 12-well plate. HDR efficiency was assessed by flow cytometry on day 5 post-transfection.

### Flow Cytometry and FACS

Flow cytometric analysis of ACDC cells was performed according to standard protocols using fluorophore-conjugated antibodies. The following reagents were used at a final concentration of 1 μg/mL in calcium- and magnesium-free D-PBS (Thermo, #14190144): PE-conjugated anti-human β2-microglobulin (B2M, clone 2M2, BioLegend #316306), APC-conjugated anti-human HLA-A,B,C (clone W6/32, BioLegend #311410), and STAT5-mRuby. Prior to staining, cells were washed once with D-PBS, incubated with antibodies for 20 min on ice, and then washed twice before analysis by flow cytometry.

For co-culture experiments with TnT cells, ACDC cells were specifically labeled with APC-conjugated anti-human podoplanin (PDPN, clone NC-08, BioLegend #337022) to distinguish them from TnT cells. T cell activation was assessed by measuring PE-Cy7-conjugated anti-human CD69 (clone FN50, BioLegend #310912) expression and NFAT-GFP reporter activity.

For soluble TCR (sTCR) binding assays, ACDC cells were incubated on ice for 20 minutes with sTCR-Fc fusion proteins, followed by two washes with calcium- and magnesium-free D-PBS. Cells were then stained with PE-conjugated anti-human IgG Fc antibody (clone M1310G05, BioLegend #410708) for 20 minutes on ice. Soluble TCRs, including TCR_MAG-IC3_-Fc and Kimmtrak-Fc, were used at final concentrations of 12.5 nM.

Flow cytometry data were acquired using BD LSRFortessa or Beckman Coulter CytoFLEX instruments. Fluorescence-activated cell sorting (FACS) was performed using BD FACSAria III or BD FACSAria Fusion instruments. Single-cell sorts were deposited into 96-well flat-bottom plates containing conditioned medium and cultured for 2–3 weeks prior to downstream characterization.

### Genotyping and transfectants

Genomic DNA was extracted from 1 × 10^6^ cells using the PureLink® Genomic DNA Kit (Invitrogen, #K182001) according to the manufacturer’s protocol. A total of 5 μL of purified genomic DNA was used as the template for 25 μL PCR reactions, performed using Q5® Hot Start High-Fidelity 2X Master Mix (NEB, #M0494S) and target-specific primers (see Supplementary Table 6). PCR products were analyzed by gel electrophoresis and, when necessary, verified by Sanger sequencing to confirm genomic edits.

### Synthetic peptides and peptide pulsing

Peptides and peptide libraries (i.e., peptide scanning libraries and potential off-target libraries) were synthesized through custom peptide synthesis (Genscript) and reconstituted in dimethyl sulfoxide (DMSO) at a concentration of 10 mg/mL. For long-term storage, peptides were maintained at -80°C. For peptide pulsing, ACDC cells were harvested and subjected to two washes with serum-free RPMI 1640 to remove residual serum components. Peptides were diluted to a final concentration of 50 μg/mL in serum-free RPMI 1640, and used to resuspend ACDC cells at a density of 1×10^6^ cells/mL. The peptide-cell suspension was incubated at 37°C with 5% CO₂ for 90 minutes, followed by a single wash with SF-RPMI to remove unbound peptides. Cells were then resuspended in complete culture media and subsequently introduced into co-culture assays as described in the following section.

### Co-culture assays for functional validation

For co-culture assays, TnT-TCR cells were harvested at a density of ∼1 × 10⁶ cells/mL, pelleted by centrifugation, and resuspended in fresh complete medium at the same concentration. A total of 1 × 10⁵ TnT-TCR cells (100 μL) were seeded into U-bottom 96-well plates. As a positive control, TnT cells were stimulated overnight with 1× eBioscience Cell Stimulation Cocktail (81 nM PMA, 1.34 μM ionomycin; Thermo Fisher, #00497093).

ACDC cells—either stably expressing antigens or peptide-pulsed—were adjusted to 1 × 10⁶ cells/mL in complete medium, and 5 × 10⁴ cells (50 μL) were added per well. Anti-human CD28 antibody (clone CD28.2, BioLegend, #302933) was included in all wells, including negative controls, at a final concentration of 1 μg/mL to provide co-stimulation. Plates were incubated overnight at 37 °C in a 5% CO₂ atmosphere.

The following day, cell activation was assessed by flow cytometry based on NFAT-GFP reporter expression and CD69 upregulation in TnT-TCR cells. In parallel, IL-2 activation in ACDC cells was evaluated via STAR5-mRuby reporter expression. TnT-TCR cells were distinguished from ACDC cells using APC-conjugated anti-human podoplanin (PDPN) staining during flow cytometric analysis.

### Generation of genomically-encoded peptide antigen Libraries

Antigen libraries were generated using single-stranded oligonucleotides (ssODNs, oPools from IDT) and assembled into HDR templates for CRISPR-mediated integration into ACDC-HLA cells. For the TCR_MAG-IC3_/MAGE-A3 library, 84 distinct ssODNs were designed, each encoding a unique combination of three NNK codon substitutions across the wild-type peptide sequence. For the Kimmtrak/gp100 library, a total of 120 ssODNs were used, consisting of 84 ssODNs with unique combinations of three NNK codons and an additional 36 ssODNs with unique combinations of two NNK codon substitutions. The RH 5k and MLP 5k libraries (Supplementary table 8 and 9) were constructed using ssODNs encoding 5,000 individually selected peptides derived from reverse Hamming (RH)-based and machine learning (MLP)-based off-target prediction models, respectively.

Each ssODN was flanked by BsaI recognition sites and approximately 20-nucleotide overhangs to facilitate second-strand synthesis by 5 cycles of PCR. Full-length peptide-encoding gene segments were assembled using the NEBridge® Golden Gate Assembly Kit (NEB, #E1602), with individual oligonucleotide fragments cloned into a custom entry vector derived from pYTK095 (Addgene #65202). This vector was engineered to contain both upstream and downstream homology arms for genomic integration, as well as a CMV promoter driving expression of β2-microglobulin (B2M).

Assembly reactions were performed using a 2:1 molar ratio of insert to vector (120 fmol vector), with 2 μL of NEB Golden Gate Enzyme Mix. The following thermal cycling protocol was used: 42 °C for 5 min and 16 °C for 5 min, repeated for 30 cycles, followed by a final incubation at 60 °C for 5 min. Assembled products were amplified by PCR using primer pair KH241/KH240 (see Supplementary Table 6), and purified using the DNA Clean & Concentrator-25 kit (Zymo Research, #D4005) to generate the final HDR templates.

### Antigen library screening and selections

Antigen library HDR templates (2 μg) and GFP-targeting gRNA (0.2 nmol) were electroporated into 5 × 10⁶ ACDC-B2MKO cells per reaction using the 4D-Nucleofector device (Lonza) with the SF Cell Line Kit (Lonza, #V4XC-2024). A total of 2 × 10⁷ cells were transfected across multiple reactions. On day 5 post-transfection, ACDC cells with restored HLA and B2M surface expression and loss of GFP expression were enriched by two rounds of bulk sorting. The resulting population, referred to as ACDC-Ag cells, was expanded for one week and subsequently used for downstream assays, including staining with soluble TCRs or co-culture with TnT-TCR cells. Cells displaying soluble TCR binding and those activated via IL-2 signaling, as indicated by STAR5-mRuby expression, were isolated by FACS for further analysis.

### Soluble TCR Expression and Purification

TCR fragments were re-formatted for soluble expression, incorporating an additional interchain disulphide bond by introducing two mutations: T166C in TRAC and S173C in TRBC (*28*). The constant TCR regions were truncated at C213 (TRAC) and C247 (TRBC1). TCR constructs were synthesized (Twist Bioscience) and cloned into the mammalian expression vector pTwist_BetaGlobin_CMV_WPRE (Twist Bioscience). Both TCRα and TCRβ chains were encoded on a single plasmid, separated by a P2A self-cleaving peptide, with either 6×His tag fused to the N-terminus of the TCRβ chain or an IgG1-Fc tag fused to the C-terminus of the TCRα to enable purification. Soluble TCRs were expressed in suspension-adapted HEK293 cells (Expi293, Thermo Fisher) transfected with the plasmids, and secreted TCRs were purified from the culture supernatant using either a 5 mL HisTrap Excel column (Cytiva, #17371206) or a 5 mL HiTrap MabSelect column (Cytiva, #28408256) on an ÄKTA Pure system (Cytiva). For His-tagged proteins, after washing with 20 mM sodium phosphate, 0.5 M NaCl, and 25 mM imidazole, TCRs were eluted in a single step using 20 mM sodium phosphate, 0.5 M NaCl, and 500 mM imidazole. For Fc-tagged proteins, after washing with 1x PBS, Fc-tagged TCRs were eluted in a single step using a 0.1M glycine buffer (pH 3.0) and neutralized with 1M Tris (pH 9.0). Purified proteins were buffer-exchanged into PBS (pH 7.5).

### Library Preparation and Sequencing of Antigen Library Amplicons

Genomic DNA was extracted from ACDC-Ag cells following selection, and peptide-encoding regions were amplified by genomic PCR using primer pair KH185/KH197. To minimize amplification bias, PCR was limited to 25 cycles. Amplicons were column-purified, and 50 ng of purified product was used for adapter ligation with KAPA UDI adapters (Roche, #08861919702) using the KAPA Hyper PCR-Free Kit (Roche, #07962371001). Size selection was performed using AMPure XP beads (Beckman, #A63881) to remove undesired fragments.

Adapter-ligated libraries were quantified by qPCR using the Roche LightCycler to ensure successful ligation and enable accurate normalization of library concentrations. Final pooled libraries were sequenced on an Illumina MiSeq platform using paired-end sequencing (2 × 151 bp) with the MiSeq Reagent Kit v2 or, alternatively, on a NovaSeq system using the SP Reagent Kit.

### Preprocessing of deep sequencing data

Sequencing reads were paired, quality trimmed and merged using the BBTools suite(BBMap 2022) with a quality threshold of qphred R>25. Sequences encoding presented peptides were then extracted using custom R scripts, followed by translation to aa sequences. Read counts per sequence were calculated and singletons (read count = 1) were discarded. Sequencing datasets used for training machine and deep learning models were created by combining the datasets from binding (positive) and non-binding (negative) sequential sorts. Sequences present in both populations were assigned a positive label, as some portion of the peptide still maintained binding to the TCR in order to be present in the positively sorted set.

Binding scores for heatmaps shown in (Fig. 3F) were created by calculating relative enrichment of ED1 sequences between the unsorted library and the sorted library. Relative enrichment was calculated based on AA frequencies calculated from total aa counts per position in each dataset. During relative enrichment calculation, a pseudocount of 1 was added to those peptides that were unsequenced in the original unenriched library, but not to the sorted library.

### Training and testing machine and deep learning models

All machine learning code and models were built in Python (3.10.4). For data processing and visualization, numpy (1.23.3), pandas (1.4.4), matplotlib (3.5.3) and seaborn (0.12.0) packages were used. Baseline benchmarking models were built using Scikit-Learn (1.0.2), while Keras (2.9.0) and Tensorflow (2.9.1) were used to build deep learning models. To prevent data leakage, all 9-mers present in the human peptidome derived dataset were removed from the input dataset before use for model training and evaluation.

During baseline model testing, 10% of the full dataset was split into a held-out test set to be used to evaluate the performance of all models, including the MLP ensembles. Baseline models were trained on 5 different splits of the remaining 90% training data to obtain final model performances. The training dataset was balanced during training, while validation and test sets remained unbalanced to appropriately evaluate MCC, precision and recall scores on imbalanced data. Dataset balancing was performed using the SMOTE oversampling strategy from the imbalance-learn package (0.11.0) for models from Scikit-Learn, while a custom data sampler was created in Tensorflow for use with deep learning MLP models.

Peptide library sequences were one-hot encoded into a matrix, and then flattened into a one-dimensional vector, prior to being used as inputs into the models. During training of classical machine learning models (Logistic Regression, SGD and XGBoost), 30 rounds of random hyperparameter search were performed using RandomSearchCV, and the best model scores were used in comparative analysis. The base MLP model consisted of two dense layers with 32 neurons each, trained with the Adam Optimization algorithm, with default parameters.

To train the final ensemble of 200 MLP deep learning models, exhaustive hyperparameter search was performed to optimize performance through the hyperparameters listed in (Supplementary Table 4). The final ensemble models were trained using the full dataset, with different random 90/10 training-validation splits to make sure each model learned slightly different parameters of the full peptide sequence space, however no test set was used for final evaluation, as to give the model as much data as possible during training.

### Preparation of human proteome-derived 9mer dataset

The Human Proteome dataset from UniProt (Proteome ID UP000005640) was accessed in October 2022, and cut into all potential 9mers using a custom script with a 1-residue sliding window. Of note, the dataset only contains the canonical sequence of each protein, and does not contain alternative isoforms. Following this, peptides containing stop codons (*) or unnatural amino acids were removed from the dataset. Finally, each 9-mer sequence was predicted for pMHC HLA-I presentation using the NetMHCPan 4.1 prediction tool available on IEDB. (https://nextgen-tools.iedb.org/pipeline?tool=tc1)

### Predictions made with ensemble deep learning models

Human peptidome 9-mers were predicted for binding using an ensemble of trained models by soft (or weighted) voting. For a given peptide sequence, each model returns raw probabilities of binding between 0 and 1. The average of all probabilities from the full 200 MLP ensemble were then obtained as the final binding probability of each peptide.

### Rank Score calculation to select top enriched human peptides from sorted libraries

Given that multiple peptides were observed in both the positive and negatively selected populations, it was important to consider the relative enrichment of each peptide in the positive compared to the negative fractions, as well as the absolute enrichment of each peptide compared to the parental, unsorted libraries. Thus we computed a ranking-based score that integrates both relative enrichment (ratio rank) and absolute enrichment (enrichment rank), to select peptides highly enriched in the positive selection condition while simultaneously being depleted in the negative selection condition receive the highest rankings.

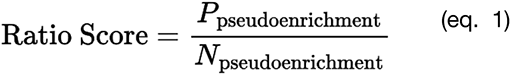

First, the enrichment of each peptide in the positive and negative populations was calculated relative to the original, unsorted parental library. In cases where the peptide was not observed in either population, it was assigned lowest observed enrichment value across the dataset to avoid undefined ratios during calculation. Then, a ratio of positive over negative enrichment was calculated to obtain a “ratio score” (eq. 1), which was then ranked in descending order to obtain a “ratio rank”. At the same time, an “enrichment rank” was obtained for each peptide based on their enrichment values in the positive dataset. The final “Rank Score” was computed as the average of the two rankings (eq. 2).

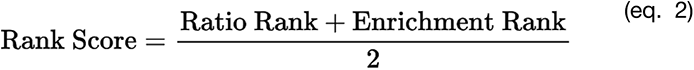

### Reverse Hamming scoring of peptides

RH scoring of peptides were adapted from Gee et al. (Gee et al. 2018) Briefly, for each target human 9-mer peptide, the Hamming distance to every synthetic peptide in the positively selected dataset was calculated and then subtracted from the total peptide length (which is 9). This score represents the similarity of each human peptide to the positively sorted set of synthetic peptides. To obtain the final RH score for each human 9-mer peptide, the reverse Hamming distance to each synthetic library peptide was weighted. These weights were determined by the rank of the synthetic peptide’s count within the positively sorted dataset. Specifically, the counts of synthetic peptides from the relevant sort round in the positive library data were ranked in ascending order. These ranks were then normalized by dividing by the maximum rank to obtain the weights. Thus, those with the highest count would have the greatest weight, and vice versa. Finally, the weighted reverse Hamming distances were summed and then divided by the total number of synthetic peptides in the positive dataset to yield the final RH score for each human peptide.

### AlphaFold3 structural modeling and comparative analysis of TCR–pMHC interactions

TCR–HLA–peptide complexes were modeled as multimers using AlphaFold3, where the HLA, B2M, TCR α- and β-chains, and the peptide sequence were provided as separate input chains. For each complex, 10 independent seeds were used to generate predictions. From each seed, the top five diffusion-ranked models were selected, resulting in 50 structures per complex for downstream analysis.

TCR binding site residues were defined as all residues in the TCR α- and β-chains that are within 5 Å of any peptide atom. Confidence of the predicted TCR–peptide interaction was quantified by calculating the mean per-atom pLDDT across all atoms of the identified binding site residues.

All predictions were performed on an NVIDIA A100 GPU.

---

**Fig. S1.**
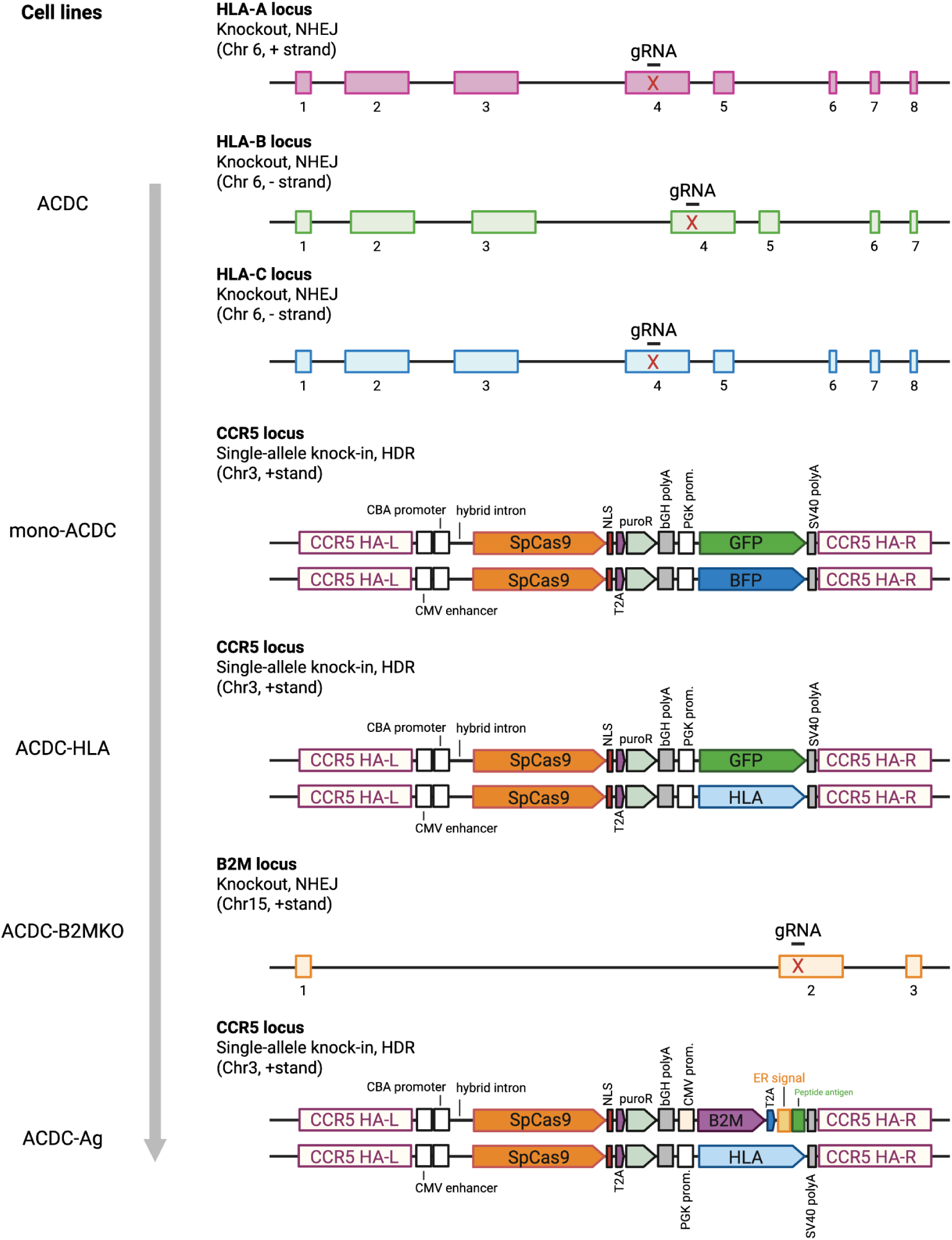
Development of the ACDC cell lines through multi-step CRISPR-Cas9 genome editing Schematic representation of the genome editing steps performed for generating the ACDC cell lines. GFP, green fluorescent protein; BFP, blue fluorescent protein; gRNA, guide RNA; bGH polyA, bovine growth hormone polyadenylation signal; Puro, puromycin; HA, homology arm; HDR, homology-directed repair; NHEJ, non-homologous end joining; NLS, nuclear localization signal; T2A, 2A peptide from Thosea asigna virus capsid protein; PGK prom, promoter of phosphoglycerate kinase; Puro, puromycin; ER signal, endoplasmic reticulum signal sequence; CBA, chicken β-actin promoter; SpCas9, Streptococcus pyogenes Cas9; SV40 polyA, simian virus 40 polyadenylation signal; B2M, Beta-2 microglobulin.

**Fig. S2.**
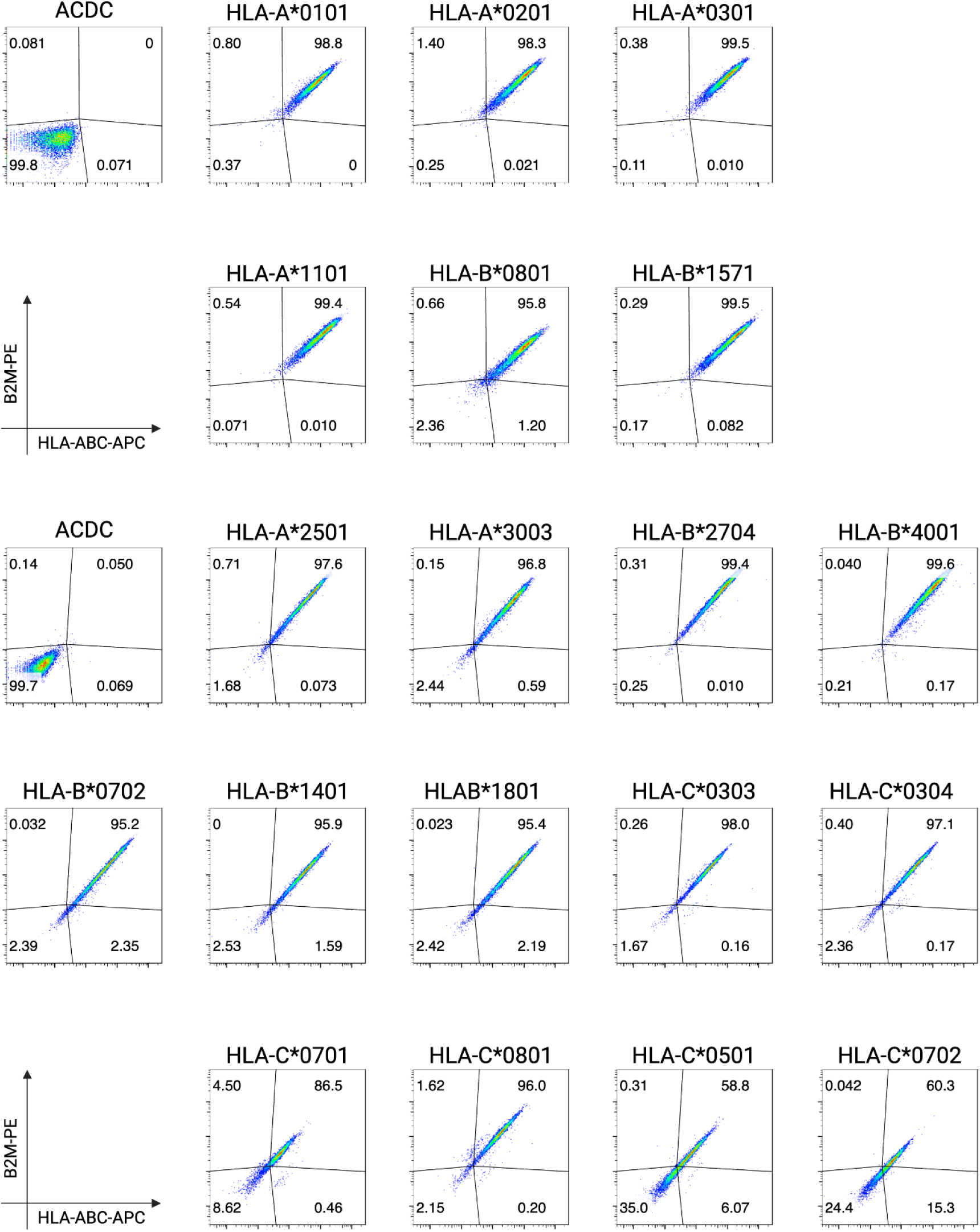
Generation of ACDC-HLA cell lines across 18 distinct HLA class I alleles. Flow cytometry analysis of ACDC cells transfected with HDR templates encoding HLA class I transgenes. Successful genomic integration of HLA alleles was assessed by restoration of HLA and B2M surface expression, as detected by HLA-ABC–APC and B2M–PE antibody staining, respectively. Each plot represents a different HLA class I allele integrated at the CCR5 locus. ACDC cells lacking transgene expression serve as negative controls.

**Fig. S3.**
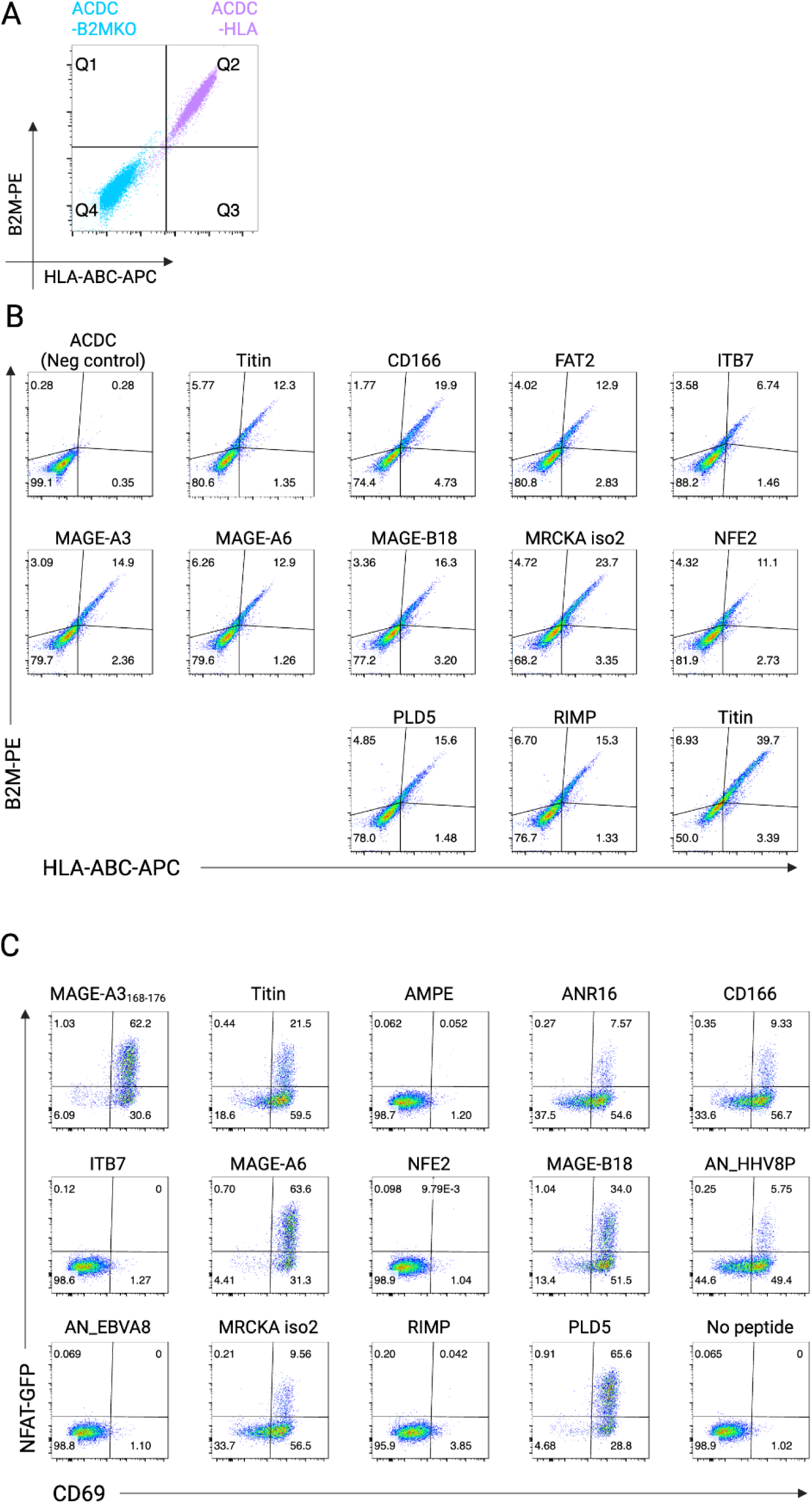
Validation of antigen knock-in and TCR-dependent activation in ACDC-Ag cells. (A) Flow cytometry plots of ACDC-B2MKO and ACDC-HLA cells showing efficient knockout of B2M expression following CRISPR targeting. (B) Flow cytometry analysis of ACDC-B2MKO cells five days post-transfection shows efficient reconstitution of B2M and 13 different genomically encoded peptide antigens expression following CRISPR-mediated HDR. (C) Functional screening of the same antigen panel using TnT cells expressing the affinity-enhanced TCR_a3a_ specific for MAGE-A3. Co-cultures were assessed for T cell activation via NFAT-GFP and CD69 expression, indicating specific activation by cognate pHLA ligands presented on ACDC-Ag cells. Peptide-negative ACDC-HLA cells served as negative controls.

**Fig. S4.**
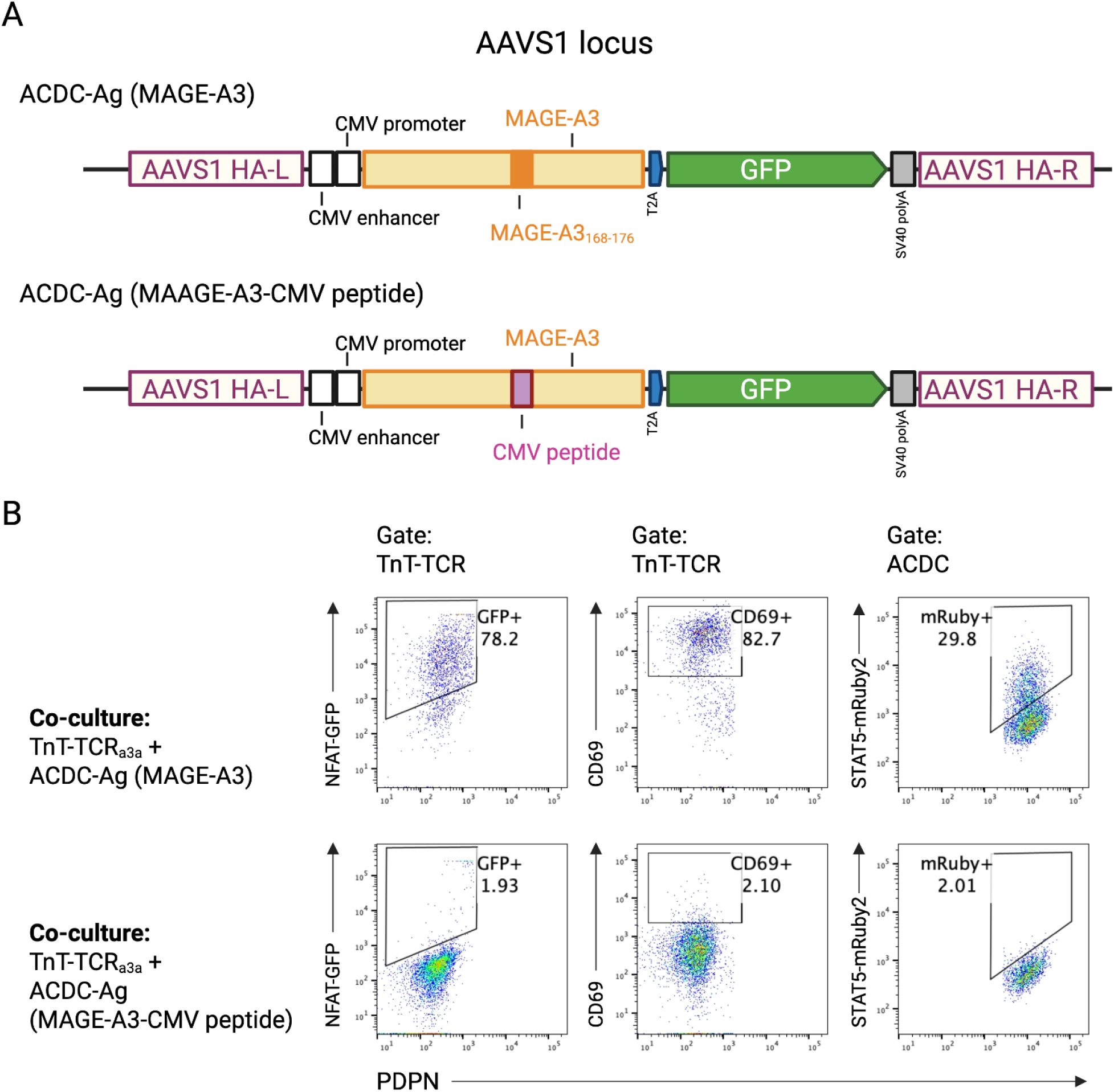
Functional presentation of full-length MAGE-A3 antigen in ACDC-Ag cells with TnT-TCR_a3a_. (A) Schematic of the HDR donor constructs for genomic integration of full-length MAGE-A3 or MAGE-A3 fused to a CMV-derived peptide into the AAVS1 locus of ACDC cells. Constructs include a CMV promoter, MAGE-A3 coding sequence, and a T2A-linked GFP reporter. (B) Flow cytometry analysis of co-cultures between TnT-TCR_a3a_ cells and ACDC-Ag cells expressing full-length MAGE-A3 (top row) or MAGE-A3-CMV peptide fusion (bottom row). Activation markers include NFAT-GFP and CD69 expression in TnT-TCR cells, and mRuby2 fluorescence in ACDC cells. Robust activation was observed only in co-cultures with full-length MAGE-A3, confirming endogenous processing and presentation of the MAGE-A3 peptide antigen.

**Fig. S5.**
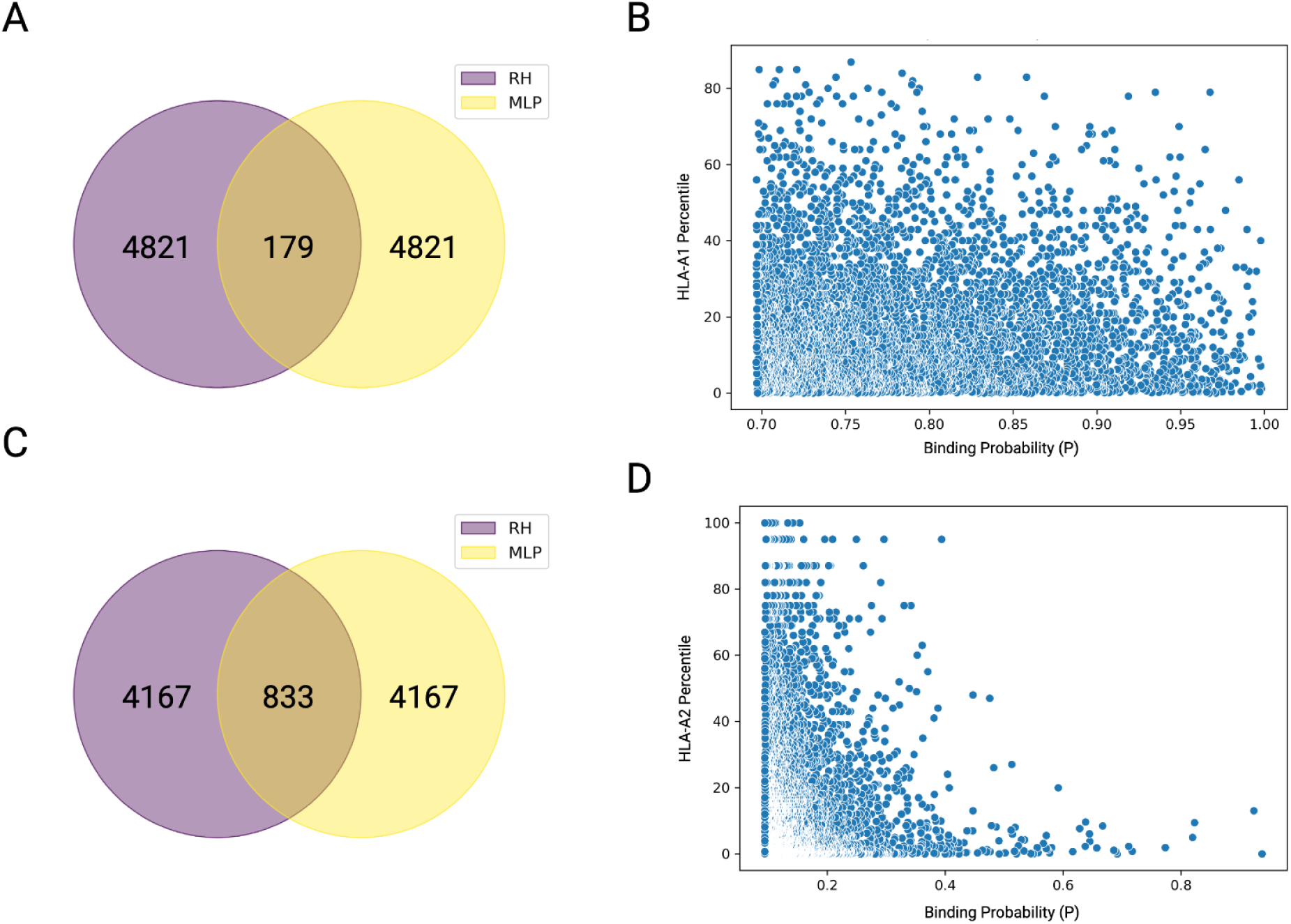
Comparison of RH and MLP predictions for TCR_MAG-IC3_ and Kimmtrak across the human proteome. (A) Venn diagram showing the overlap of the top 5,000 predicted peptide binders to TCR_MAG-IC3_ identified by RH scoring (purple) and MLP ensemble models (yellow). Only 179 peptides were shared between the two methods. (B) Scatter plot showing MLP ensemble binding probability scores versus NetMHCpan-predicted percentile ranks for HLA-A*01:01 presentation of the top 5,000 TCR_MAG-IC3_ peptides. (C) Venn diagram showing the overlap of the top 5,000 predicted peptide binders to Kimmtrak from RH scoring (purple) and MLP ensemble models (yellow). A higher overlap of 833 peptides was observed between the two methods. (D) Scatter plot of MLP ensemble binding scores versus NetMHCpan-predicted percentile ranks for HLA-A*02:01 presentation of the top 5,000 Kimmtrak peptides.

**Fig. S6.**
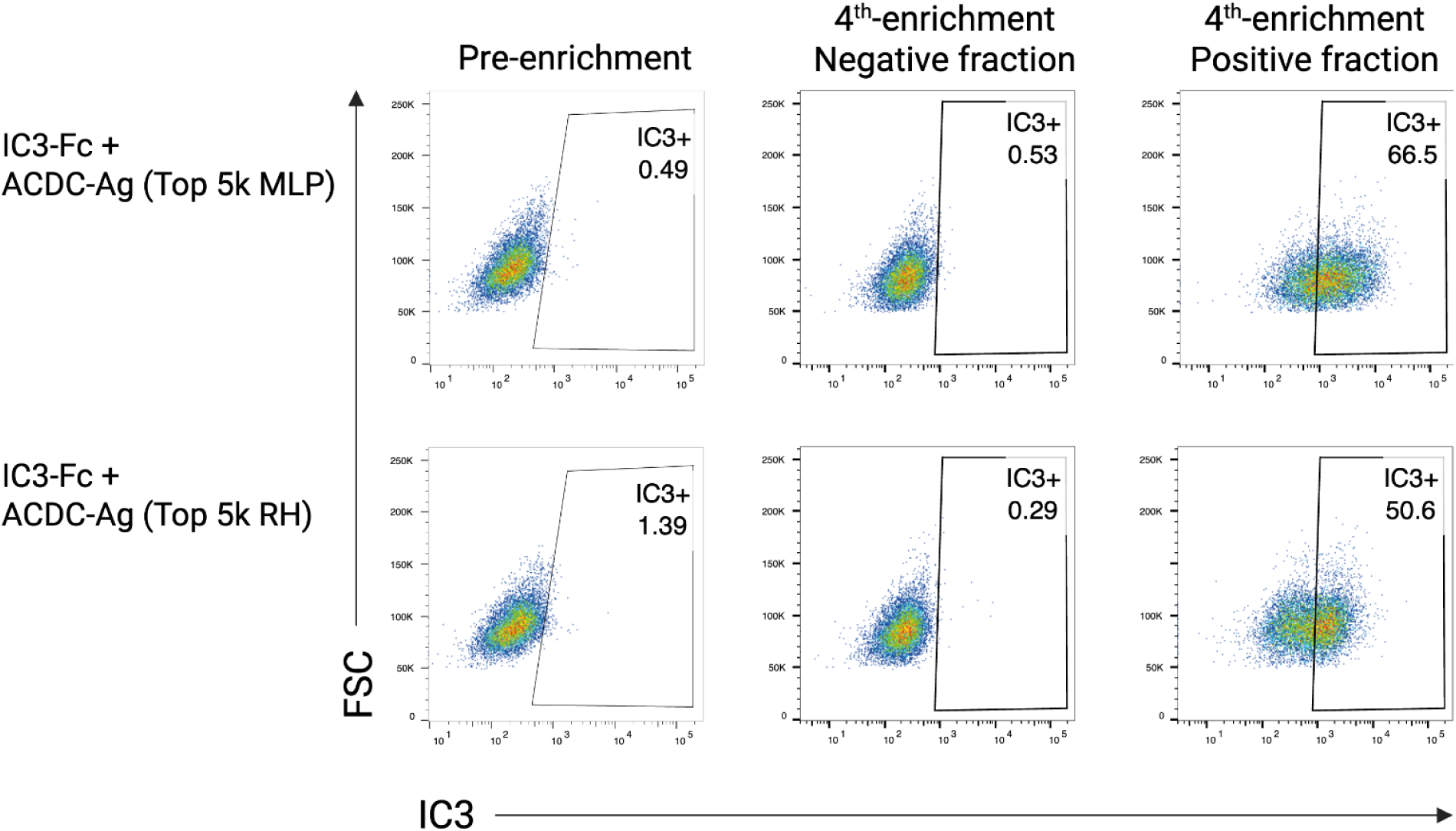
Pooled functional screening of the predicted top 5,000 off-target peptides using ACDC-Ag cells. Representative flow cytometry plots showing enrichment of TCR_MAG-IC3_-Fc–positive ACDC-Ag cells expressing the top 5,000 peptide candidates predicted by either MLP ensemble (top row) or reverse Hamming (RH) scoring (bottom row). Left: pre-enrichment populations showing baseline TCR_MAG-IC3_-Fc staining. Middle: fourth-round enrichment, TCR_MAG-IC3_-Fc negative-sorted fraction. Right: fourth-round enrichment, TCR_MAG-IC3_-Fc positive fraction. Both libraries yielded substantial enrichment of TCR_MAG-IC3_-Fc+ cells, with final positivity reaching 66.5% (MLP) and 50.6% (RH), respectively.

**Fig. S7.**
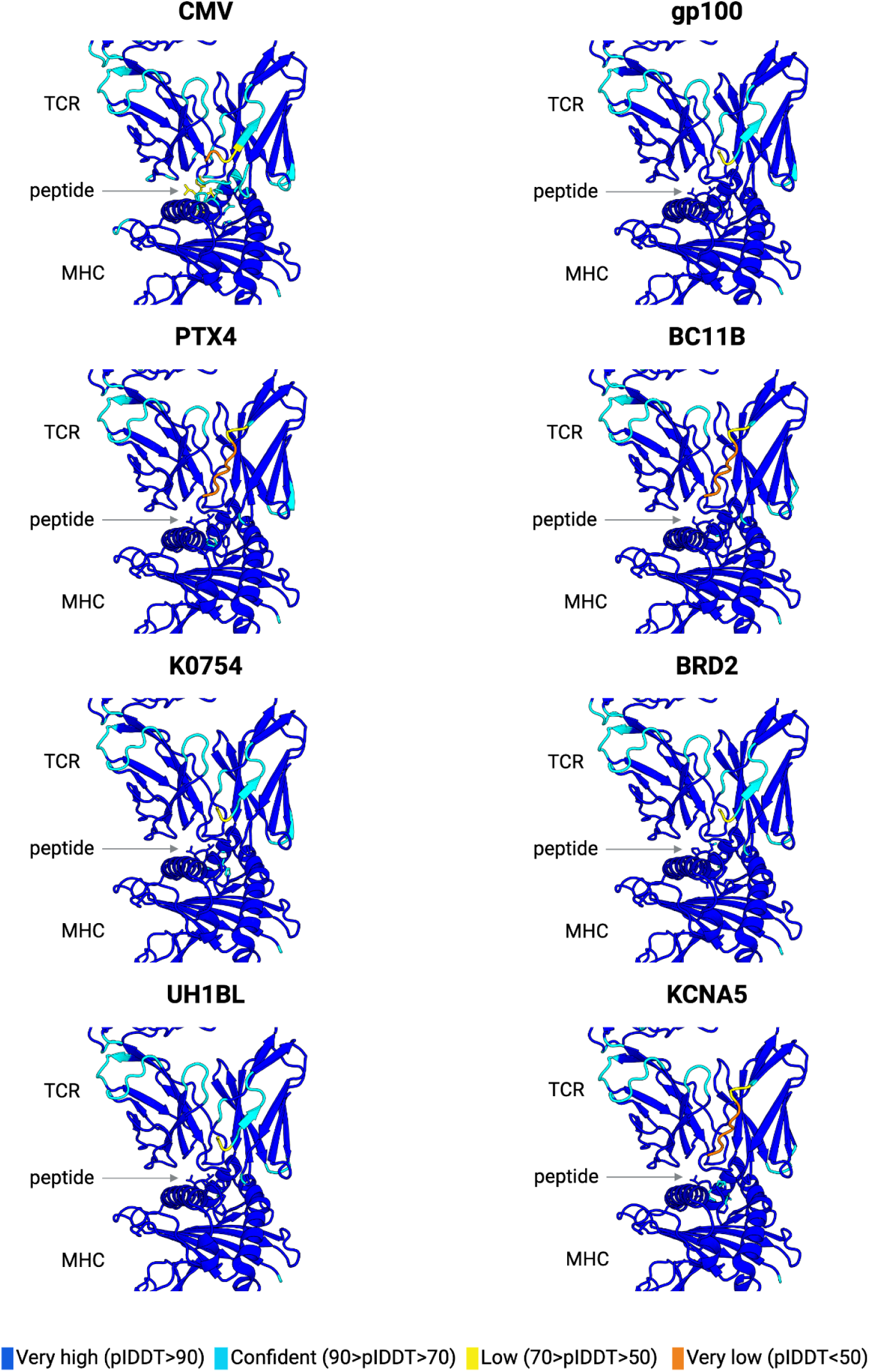
AlphaFold3 structural predictions of Kimmtrak bound to gp100, CMV and off-target peptides–HLA-A*02:01 complexes. Structural models of the Kimmtrak bound to gp100 and six validated off-target peptides (PTX4, BC11B, K0754, BRD2, UH1BL, KCNA5) presented by HLA-A*02:01, alongside the non-binding CMVpp50 peptide. Structures are colored by per-residue predicted IDDT (pIDDT) confidence scores: very high (blue, pIDDT > 90), confident (cyan, 90 > pIDDT > 70), low (yellow, 70 > pIDDT > 50), and very low (orange, pIDDT < 50).

**Table S1.** Sequencing Coverage and Diversity of the 3NNK Library of MAGE-A3.

| Library subset | Number of mutations | Theoretical diversity | Observed Unique Sequences | Coverage |
| --- | --- | --- | --- | --- |
| 1NNK | 1 | 171 | 171 | 100% |
| 2NNK | 2 | 12,996 | 7,051 | 54.26% |
| 3NNK | 3 | 576,156 | 41,524 | 7.20% |
| Total |  | 589,323 | 48,746 | 8.27% |

**Table S2.** Sequencing Coverage and Diversity of the 3NNK Library of gp100.

| Library subset | Number of mutations | Theoretical diversity | Observed Unique Sequences | Coverage |
| --- | --- | --- | --- | --- |
| 1NNK | 1 | 171 | 171 | 100% |
| 2NNK | 2 | 12,996 | 12,401 | 95.42% |
| 3NNK | 3 | 576,156 | 101,050 | 17.54% |
| Total |  | 589,323 | 113,622 | 19.28% |

**Table S3.** Performance Metrics of Machine Learning Classifiers for TCR Off-Target Prediction

| Base model | binary accuracy | f1 score | binary mcc | precision | recall |
| --- | --- | --- | --- | --- | --- |
| Logreg | 0.743 | 0.611 | 0.428 | 0.555 | 0.680 |
| SGD | 0.753 | 0.614 | 0.436 | 0.569 | 0.666 |
| XGBoost | 0.843 | 0.693 | 0.606 | 0.823 | 0.601 |
| MLP | 0.798 | 0.662 | 0.519 | 0.661 | 0.666 |
| MLP Ensemble | 0.850 | 0.753 | 0.646 | 0.773 | 0.733 |

**Table S4.** Final Hyperparameter Optimization for MLP Models

| Parameter | Range | TCR<sub>MAG-IC3</sub><br>MLP<br>Final | KIMMTRAK MLP<br>Final |
| --- | --- | --- | --- |
| Learning Rate | 0.01 - 0.0001 | 0.0001 | 0.005 |
| Optimizer | Adam, SGD | Adam | Adam |
| Minority Ratio (dataset balance) | 0.1 - 0.5 | 0.5 | 0.5 |
| Training Epochs | 15 - 75 | 25 | 20 |
| # Dense Layers | 1-3 | 1 | 2 |
| # Dense Dimensions | 32 - 512 | 96 | 96, 128 |
| Dense Dropout | 0 - 0.5 | 0.2 | 0.2 |

**Table S5.**
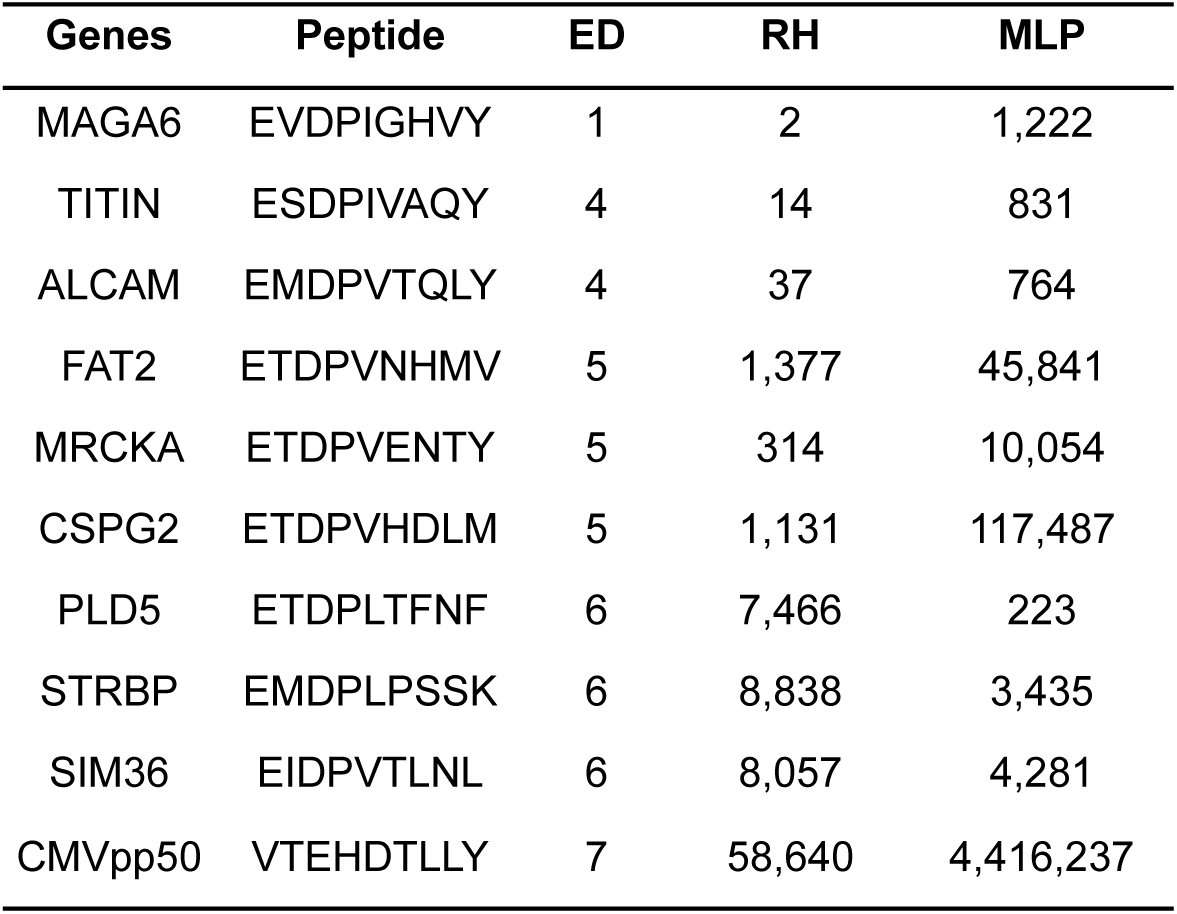
Comparison of Off-Target Ranks by RH and MLP Models on the 11M Human Peptidome 9-Mers Dataset

**Table S6.**
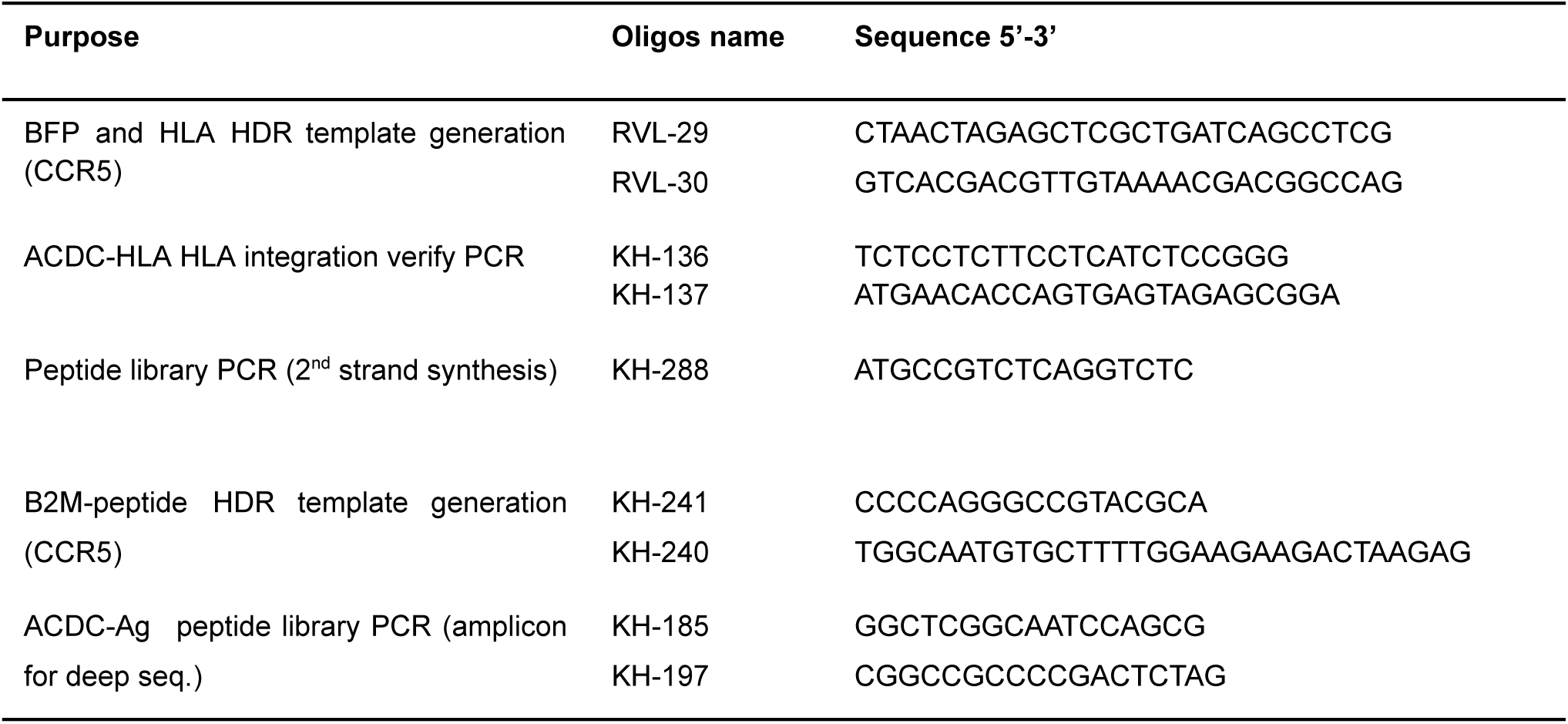
DNA oligonucleotide sequences utilized in this study

**Table S7.**
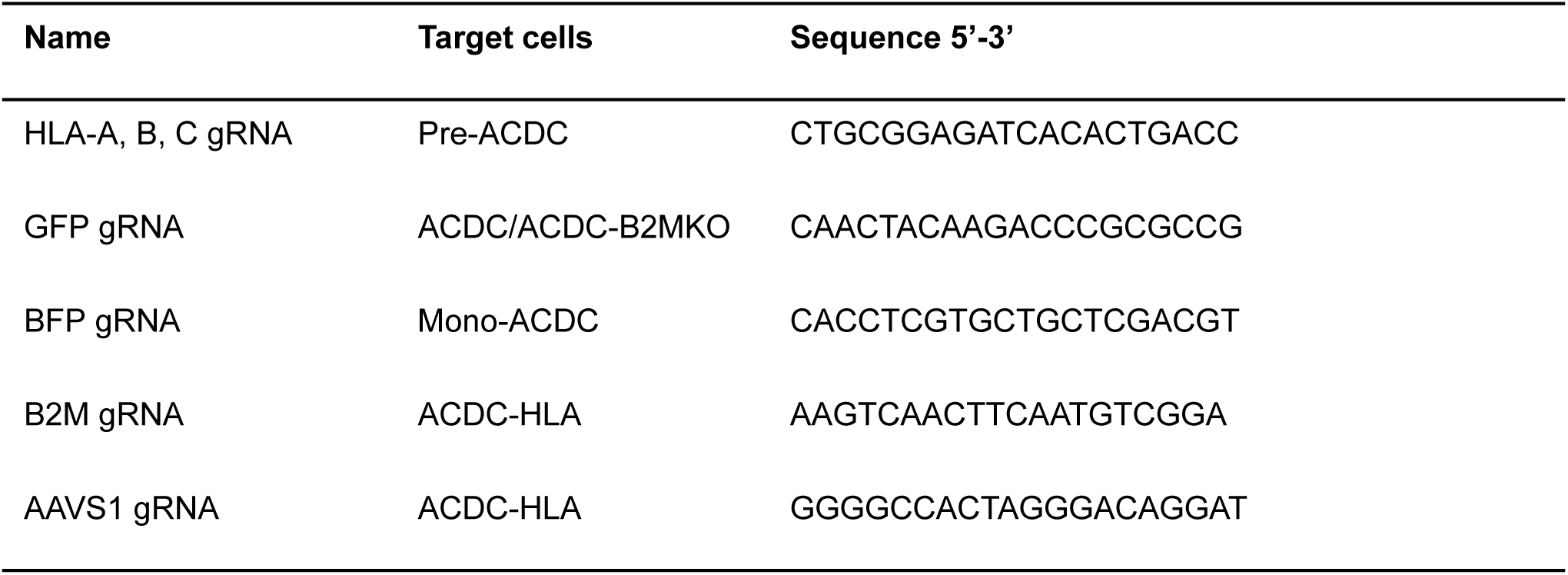
Custom Alt-R crRNA sequences (IDT) used in this study

**Table S8.** Top 5k candidate targets of TCR_MAG-IC3_ as predicted by RH and MLP

**Table S9.** Selected 95 candidate targets of TCR_MAG-IC3_ as predicted by RH and MLP

**Table S10.** Selected 40 candidate targets of Kimmtrak as predicted by RH and MLP

https://docs.google.com/spreadsheets/d/1l0c9Er1MHlb-iPaBBpyiV4XgLOZIlJHflTSiIwFXST0/edit?usp=sharing

